# Variational inference for microbiome survey data with application to global ocean data

**DOI:** 10.1101/2024.03.18.585474

**Authors:** Aditya Mishra, Jesse McNichol, Jed Fuhrman, David Blei, Christian L. Müller

**Affiliations:** Center for Computational Mathematics, Flatiron Institute, New York, NY, USA; Department of Biological Sciences, University of Southern California, LA, USA; Department of Statistics and Computer Science, Columbia Universit, NY, USA; Computational Health Center, Helmholtz Zentrum München, Munich, Germany; Department of Statistics, LMU München, Munich, Germany; Department of Statistics, University of Georgia, Athens, GA, USA

**Keywords:** Microbiome, Probabilistic model, Association learning, Variational inference, Tara ocean expedition

## Abstract

Linking sequence-derived microbial taxa abundances to host (patho-)physiology or habitat characteristics in a reproducible and interpretable manner has remained a formidable challenge for the analysis of microbiome survey data. Here, we introduce a flexible probabilistic modeling framework, VI-MIDAS (`V`ariational `I`nference for `MI`crobiome survey `DA`ta analysiS), that enables *joint* estimation of context-dependent drivers and broad patterns of associations of microbial taxon abundances from microbiome survey data. VI-MIDAS comprises mechanisms for direct coupling of taxon abundances with covariates and taxa-specific latent coupling which can incorporate spatio-temporal information *and* taxon-taxon interactions. We leverage mean-field variational inference for posterior VI-MIDAS model parameter estimation and illustrate model building and analysis using Tara Ocean Expedition survey data. Using VI-MIDAS’ latent embedding model and tools from network analysis, we show that marine microbial communities can be broadly categorized into five modules, including SAR11-, Nitrosopumilus-, and Alteromondales-dominated communities, each associated with specific environmental and spatiotemporal signatures. VI-MIDAS also finds evidence for largely positive taxon-taxon associations in SAR11 or Rhodospirillales clades, and negative associations with Alteromonadales and Flavobacteriales classes. Our results indicate that VI-MIDAS provides a powerful integrative statistical analysis framework for discovering broad patterns of associations between microbial taxa and context-specific covariate data from microbiome survey data.

## Introduction

Microbial species are an integral part of life on earth. Ecosystems, ranging from the human gut to the global ocean, harbor trillions of bacteria, archaea, viruses, and fungi that take on essential functional roles and have developed intricate ecological relationships within their respective habitat. Over the past decades, advances in amplicon and metagenomics sequencing techniques [74, 54, 52, 70] and standardized experimental and bioinformatics workflows [63, 10, 9] have enabled the large-scale collection and dissemination of microbial survey data, including those from the seminal Human Microbiome Project [69], several gut-focused surveys [28, 64, 32, 45], the Earth Microbiome Project [25], and the Tara Ocean Expedition [67]. These surveys have reached a level of maturity and complexity that ultimately allow the estimation of statistical associations between microbial abundances, typically represented as compositional counts of Amplicon Sequence Variants (ASVs) or Operational Taxonomic Units (OTUs), and habitat properties [67, 7], biogeochemical processes[29], and/or host health status [23, 48]. This, in turn, provides a starting point for deciphering and understanding the ecological and functional roles of different microbial clades in the ecosystem, nutrient and bio(geo)chemical dependencies, resource limitations of microbial growth, and the presence of ecological taxon-taxon interactions [18].

Here, we introduce an integrative probabilistic modeling framework that is specifically tailored to microbiome survey data and enables joint estimation of habitat-dependent drivers and broad associations patterns of microbial taxa abundances (see Figure 1). Our approach, termed VI-MIDAS (`V`ariational `I`nference for `MI`crobiome survey `DA`ta analysiS), models the observed taxon abundances by *simultaneously* learning taxon-specific latent representations that leverage the effects of host or environmental factors *and* taxon-taxon associations via an item-item interaction modeling *ansatz*, originally proposed for market basket analysis [61]. As such, VI-MIDAS seamlessly extends common statistical methods for microbiome data that only focus on either statistical abundance modeling [31, 39, 13, 79, 75, 49] or microbial association estimation[37, 18, 77, 57, 26]

VI-MIDAS uses the parametric structure of the Negative binomial distribution [46, 49] to account for the overdispersed nature of the amplicon count data and comprises two main model components: (i) a component that allows for full adjustment of taxon abundances from a user-defined subset of covariates and (ii) taxa-specific latent vectors that incorporate, e.g., spatio-temporal or environmental covariates *and* taxon-taxon interactions, thus providing a marginal characterization of each taxon. We resort to mean-field variational inference for parameter estimation of VI-MIDAS’ intractable posterior distribution [8], thus complementing other recent variational approaches to microbiome data modeling, such as, e.g., Poisson principal component analysis [14], microbiome dynamics modeling [24], Dirichlet Multinomial modeling [30], multi-level modeling [42], and microbiome ordination [78].

**Fig. 1.**
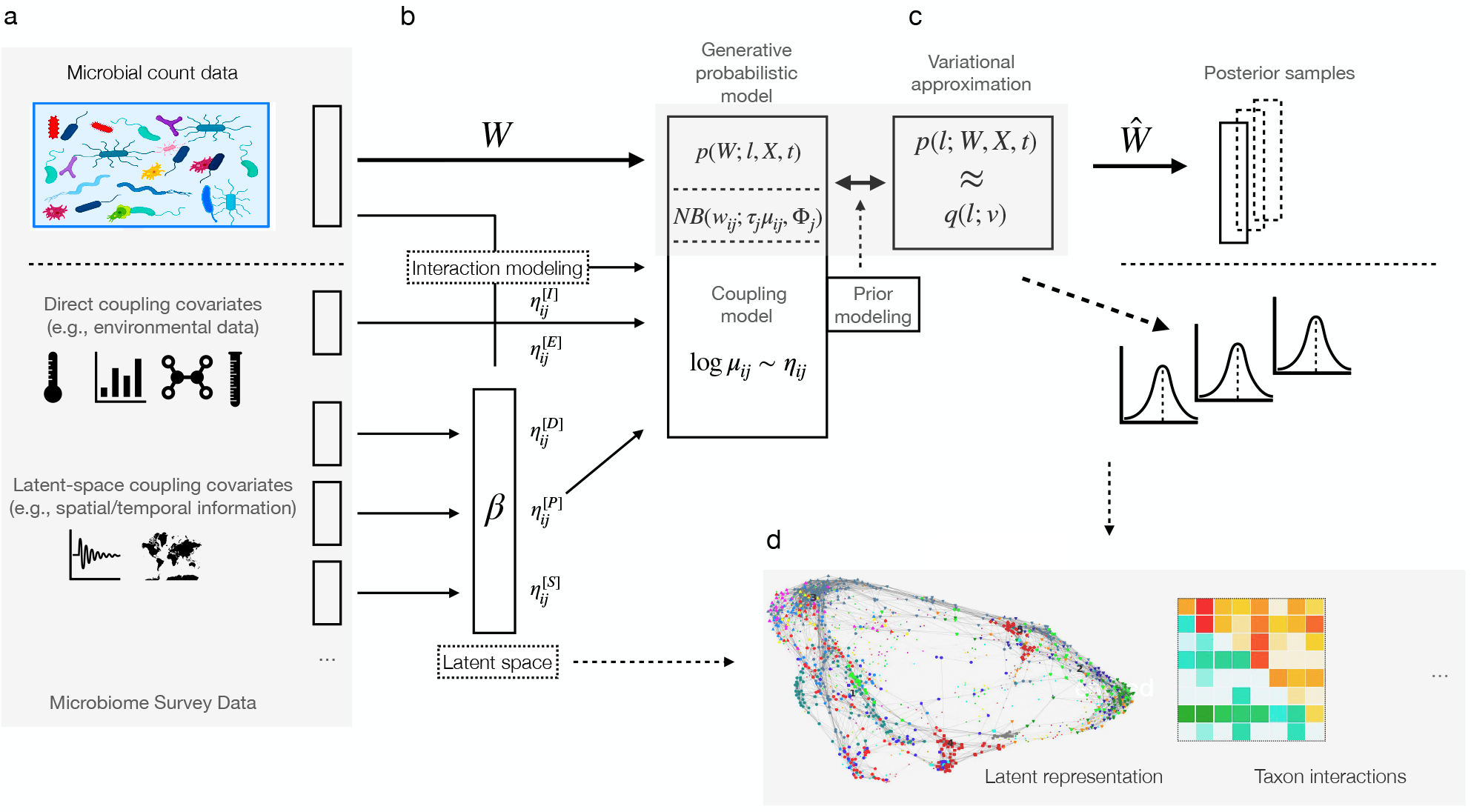
Overview of the VI-MIDAS framework. **a**. VI-MIDAS integrates microbiome survey data in form of microbial abundance data *W*, host-associated, habitat or environmental data, and spatio-temporal information. **b**. Different data sources are coupled directly *or* indirectly through a latent space ***β*** to a generative model. An additional latent space taxon interaction model is included. The generative probabilistic model (e.g., Negative Binomial (NB) model) integrates covariate data via a coupling model. **c**. Variational approximation and mean-field estimation are used for Bayesian parameter estimation, resulting in posterior microbial abundance samples 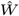 and model parameter distributions. **d**. Model components, such as estimated latent representation and taxon-taxon interactions, can be used for data understanding, visualization, and downstream analysis.

To illustrate the complete workflow of the VI-MIDAS framework, we focus on integrative analysis of global marine microbiome survey data. The ocean microbiome is of fundamental importance for life on earth, being responsible for about half of all primary production (i.e., the production of chemical energy in organic compounds) and holds enormous potential for climate remediation [50]. Several initiatives such as the Tara Oceans Project [56] and the Simons CMAP [4] provide well-structured sequencing data, biogeochemical and environmental covariate data, and satellite-derived products that are amenable to statistical analysis. Here, we re-analyze Tara expedition data ^1^, originally considered in [67] to study the structure and function of the global ocean microbiome. The expedition collected ocean water samples from 68 distinct geographical locations at varying levels of depth. We will make extensive use of this dataset to motivate and describe the details of the VI-MIDAS framework as well as the learned representations and associations of the global ocean microbiome.

We start with an overview of the Tara Oceans data under study, introduce the generative model components of VI-MIDAS, and show how different data types enter the modeling framework. We then give a high-level overview of the variational parameter estimation procedure, including the selection of VI-MIDAS’ hyperparameters, such as the choice of the priors and the dimensionality of the latent representation. Following model parameter inference, we illustrate how standard modularity analysis of VI-MIDAS’ learned latent representation of the Tara data identifies five distinct groups of microbial consortia. We analyze the inferred modules in terms of their composition of ecologically relevant clades and discuss the derived module-specific environmental and spatiotemporal signatures. Finally, we highlight the emerging interaction pattern among ecologically relevant clades and discuss the framework in the larger context of other microbiome survey data. Further methodological details are summarized in the Supplemental Material. Code for the presented VI-MIDAS workflow is available at http://github.com/amishra-stats/vi-midas) and requires minimal adjustment to analyze other microbiome survey data.

## Materials and Methods

### Tara ocean data and ecologically relevant taxa re-classification

We consider the processed Tara expedition data, as provided at http://ocean-microbiome.embl.de/companion.html. The expedition collected water samples from 68 distinct geographical locations (Figure 2b) across different depths, resulting in *n* = 139 distinct samples. Across these samples, the original data comprises microbial taxa abundances profiles of more than 35,000 bacterial taxa in form of metagenomic OTUs (mOTUs) (derived using the miTAGS framework [68]).

**Fig. 2.**
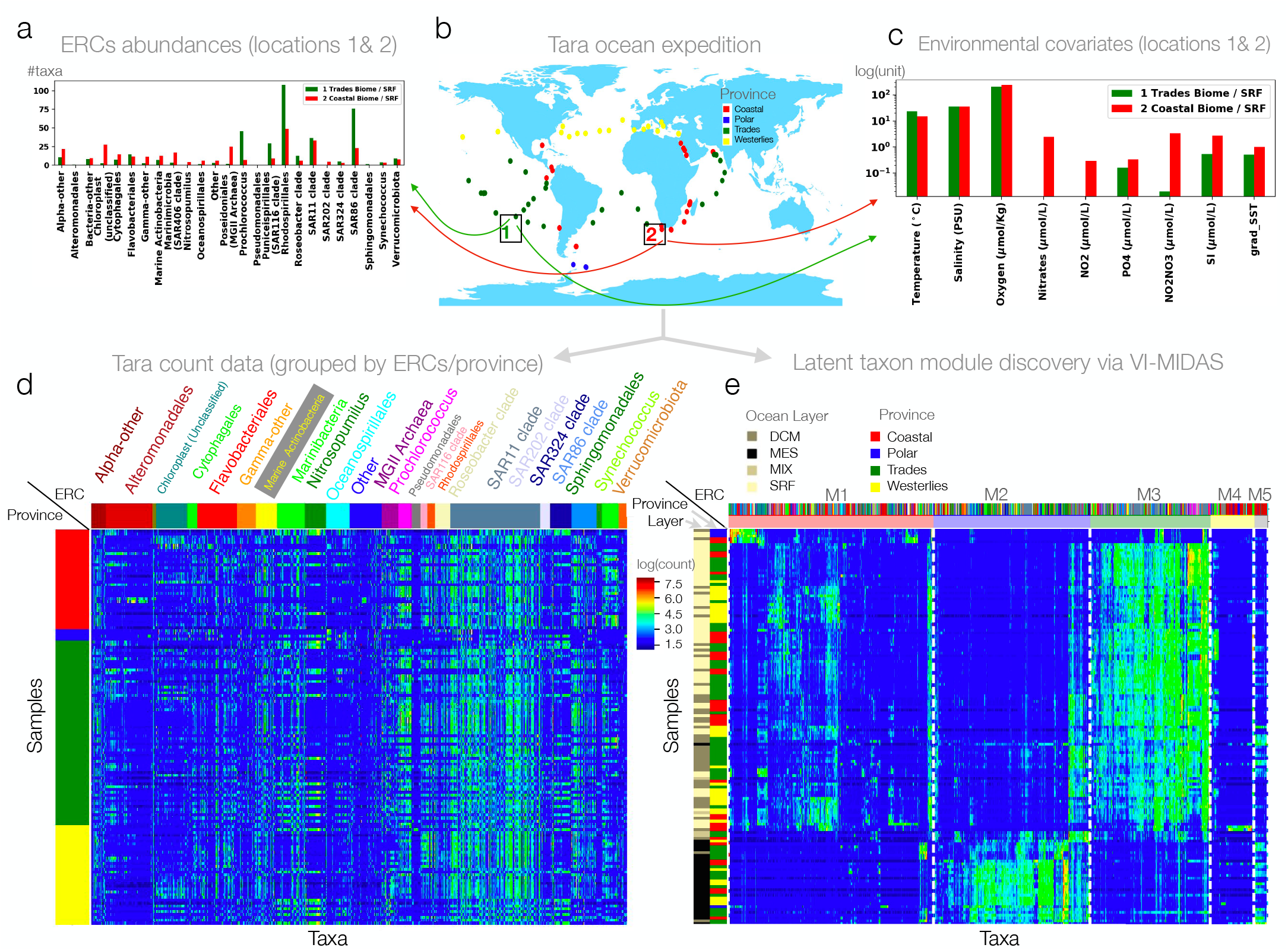
Illustration of the Tara ocean data: **a**. Taxon abundance profiles, agglomerated to expert-derived ecologically relevant classes (ERCs) for two samples (red and green, marked as 1 and 2 in Figure 2b). **b**. Tara ocean sample locations. **c**. Environmental features associated with the samples marked as 1 and 2 in Figure 2 (b); **d**. Abundance profiles log(**W** + 1) of *q* = 1379 taxa at *n* = 139 distinct locations with rows highlighting province of the sample and columns grouped by ERC. **e**. Abundance profiles clustered into five modules (M1-M5) as identified by modularity analysis of the latent space ***β*** (see Section Modularity analysis for more details). The dashed vertical lines separate the latent modules. The five microbial modules (M1-M5) comprise 524, 400, 307, 112 and 35 taxa/OTUs, respectively. The first column shows ocean depth layer, the second column the province indicator.

Here, we focus on the most abundant taxa by taking the union of all mOTUs that, in each individual sample, contribute to 40% of the total library size. This filtering allows us to cover the abundance profiles of the *q* = 1378 taxa with the most significant variability and reduces the number of excess zero counts. To account for the highly variable sequencing depth across the samples, we normalize the abundance data with respect to the lowest library size via common-sum scaling [46]. Figure 2d shows the log-transformed abundance profiles **W** ∈ ℝ^*n×q*^. Since the original taxonomic affiliations of the miTAGS are difficult to interpret, we next developed a partitioning of the selected taxa into ecologically relevant classes (ERCs). The original full taxonomy strings are too long to understand at a glance, and parsing by taxonomic level is not a good option since taxa vary widely in the depth of their annotations. For example, cyanobacteria should be annotated at the genus level or higher, but many other abundant but less described taxa do not have any taxonomic information at that level. We manually curated the data to provide a short relevant taxonomic indicator that provides a rough indicator of the ecological niche of an organism while remaining short enough to be interpreted at a glance. Some taxonomies have been altered to preserve the updated SILVA taxonomy (i.e., Betaproteobacteria is now Burkholderiales). New SILVA 138 [58] taxonomies have been used wherever possible (i.e., when the original ID was still in SILVA 138), but in cases where there was only the SILVA 108 taxonomic information, we have used our best guess. For example, if an organism had the same classification as other organisms in SILVA 108, we have often given it the same name as its counterparts in SILVA 138. We present all our findings in terms of these 29 ERCs.

Each Tara sample also contains environmental and spatiotemporal information, including geolocation, the derived Longhurst province (biome) indicator, sampling date, ocean depth information (depth from sea surface), environmental covariates, such as, e.g., sea surface temperature (SST), and biogeochemical features such as salinity, chlorophyll, nitrate, and oxygen concentration (see Figure 2c for illustration). Table 1 summarizes the measured covariates and derived spatiotemporal indicator variables that are included in the VI-MIDAS framework and their corresponding mathematical representation.

**Table 1.**
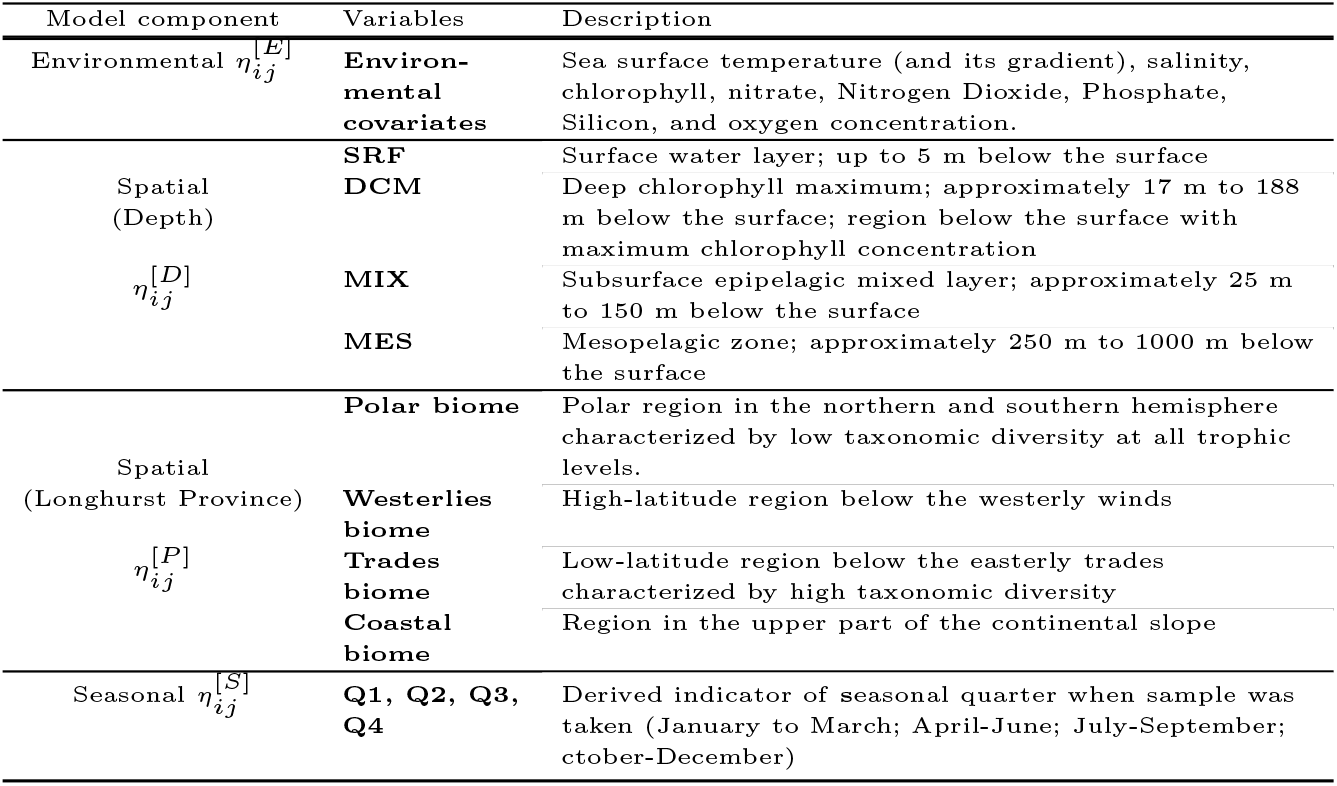
Environmental and spatiotemporal variables included in the VI-MIDAS model.

### Generative Modeling in VI-MIDAS

We seek to model the abundance profiles of *q* microbial taxa where we denote a single sample by the random variable **w** ∈ ℝ^*q*^ and the observed data from *n* samples by **W** = [*w*_*ij*_ ]_*n×q*_ ∈ ℝ^*n×q*^. For concreteness, we illustrate model building and analysis using the Tara abundance profiles (see Figure 2(d)) of *q* = 1378 taxa but the modeling strategy is applicable to any multimodal microbiome survey.

#### Distributional model

VI-MIDAS posits that the overdispersed microbial count data **W** are reasonably well modeled with the Negative Binomial distribution [11, 44, 48]. While other generative statistical modeling approaches are available, including the Dirichlet Multinomial (mixture) framework [31, 71], latent Dirichlet allocation [62], and Poisson distribution models [39, 5, 75], we found the Negative Binomial model to be an excellent choice for the Tara ocean data (see Figure S1 (b) of the Supplementary Material for the over-dispersion analysis). Using the Negative Binomial distribution with mean and dispersion parameterization [11], VI-MIDAS models the *j*th taxa in the *i*th sample as:

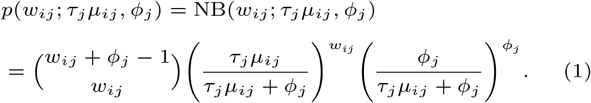

Here, the mean parameter *τ*_*j*_ *μ*_*ij*_ is the product of a taxon-specific shape parameter *τ*_*j*_ ∈ (0, 1) and the entry-specific parameter *μ*_*ij*_ ∈ ℝ^+^. The parameter *φ*_*j*_ ∈ ℝ^+^ is the taxon-specific dispersion parameter. Let us denote the dispersion and shape parameters for *q* outcomes by **Φ** = [*φ*_1_, …, *φ*_*q*_] and ***τ*** = [*τ*_1_, …, *τ*_*q*_], respectively. The shape parameter ***τ*** accounts for the disparity in abundance among microbial taxa. The generative model (1) of VI-MIDAS thus implies 𝔼(*w*_*ij*_) = *τ*_*j*_ *μ*_*ij*_ and 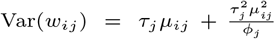. Consequently, Var(*w*_*ij*_) *>* 𝔼(*w*_*ij*_), thus making the parametric framework (1) suitable for modeling the overdispersed count data.

#### Modeling strategy and model components

One novelty in VI-MIDAS is the combination of ideas from generalized linear modeling [11] and compositional data analysis [2] to associate the microbial relative count data with spatiotemporal, environmental, and taxa information. Specifically, we model the log-transformed mean parameter ***μ*** = [*μ*_*ij*_ ]_*n×q*_ of the generative model (1) with two components, a consistent zero-aware geometric mean estimate *t*_*i*_ and a linear predictor ***η*** = [*η*_*ij*_ ]_*n×q*_ ∈ ℝ^*n×q*^ as follows:

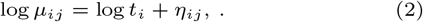

The sample-wise parameter *t*_*i*_ is estimated by a zero-aware geometric mean estimator, introduced in [16], which provides a principled approximation to the geometric means across all *n* samples in the presence of excess zeros. We detail the exact formulation of *t*_*i*_ and its approximation guarantees in Section 3.1 of the Supplementary Material. Including **O** = [log *t*_1_, …, log *t*_*n*_] as an offset term in the model is necessary since we do not have access to absolute microbial abundance data, thus requiring transforming the compositional data appropriately. The second term ***η*** effectively models centered log-ratio (clr) transformed (rather than the original count) data and is the key component to couple habitat (or host) information to the microbial abundance profiles. VI-MIDAS introduces a novel decomposition of the component ***η*** that allows the incorporation of three distinct coupling mechanisms: (i) a direct coupling term for covariates, (ii) an indirect coupling term for covariates via a latent space representation, and (iii) a latent taxon-taxon interaction term.

In our ocean application, the first component, denoted by 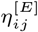, includes all relevant environmental attributes (see first row in Table 1). All spatiotemporal features, i.e., the Longhurst **P**rovince indicator, the **D**epth information, and the **S**easonal indicator (see second to last row in Table 1) are handled by the latent coupling term and are denoted by 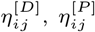, and 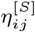, respectively. Lastly, statistical associations among co-occurring taxa are included via a latent interaction term 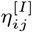, leading the following model:

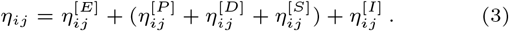

The following paragraphs detail the parametric form of each of the components, the nature of the underlying covariate data, and their biological relevance.

#### Direct coupling of environmental features

Let us denote the *p* covariates in the direct coupling term by **X** = [**x**_1_, …, **x**_*n*_]^T^ = [*x*_*ij*_ ]_*n×p*_. VI-MIDAS models the direct component for the *j*th taxa in the *i*th sample via

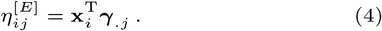

with ***γ*** = [*γ*_*ij*_ ]_*p×q*_ ∈ ℝ^*p×q*^ denoting the matrix of all coefficients. For the Tara data, we opted to model 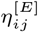 using following *p* = 9 covariates: sea surface temperature (SST) (and its gradient grad SST), salinity, chlorophyll, nitrate, nitrogen dioxide, phosphate, silicon, and oxygen concentration. All variables are mean-centered prior to incorporation into the model. In the original Tara analysis [67], temperature and oxygen have been identified as key drivers of taxonomic compositions. The VI-MIDAS analysis will allow a refined picture of the these general tendencies.

#### Latent space coupling of spatiotemporal features

VI-MIDAS offers a second mechanism for including variables of interest through latent space modeling. We denote *q* taxa-specific shared latent variables of size *k* by ***β*** = [*β*_*ij*_ ]_*k×q*_ ∈ ℝ^*k×q*^. The size factor *k* is an application-specific hyper-parameter that controls the expressiveness of the latent space. Features are then coupled to the latent space in a multiplicative fashion.

For the Tara data, we illustrate this mechanism by coupling all available spatial and temporal indicators to the latent space component. We first consider the *r* = 4 primary provinces (or biomes): polar, Westerlies, coastal, and Trades [43]. We denote the model matrix indicating the *r* distinct regions of the *n* samples by **R** = [**r**_1_, …, **r**_*n*_]^T^ ∈ ℝ^*n×r*^ and connect it to the joint latent space via the coefficient matrix ***α*** = [*α*]_*r×k*_ ∈ ℝ^*r×k*^, leading to

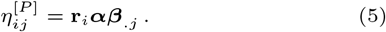

Similarly, the Tara data includes samples across *b* = 4 ocean depths: surface water (SRF), deep chlorophyll maximum (DCM), the subsurface epipelagic mixed layer (MIX), and the mesopelagic zone (MES). We denote the depth indicator matrix of the *n* samples by **D** = [**d**_1_, …, **d**_*n*_]^T^ ∈ ℝ^*n×d*^ and connect it to the joint latent space via the coefficient matrix ***δ*** = [*δ*]_*b×k*_ ∈ ℝ^*b×k*^, leading to

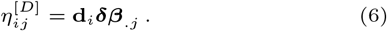

Finally, by parsing the sampling dates at the different Tara locations, we can associate a temporal indicator with each sample. Here, we group the samples into *m* = 4 seasons: the 1^*st*^ (Q1, January-March), 2^*nd*^ (Q2, April-June), 3^*rd*^ (Q3, July-September), and 4^*th*^ (Q4, October-December) yearly quarter, and construct the season indicator matrix **S** = [**s**_1_, …, **s**_*n*_]^T^ ∈ ℝ^*n×s*^. The coefficient matrix ***ϑ*** = [*ϑ*]_*m×k*_ ∈ ℝ^*m×k*^ couples **S** to the latent space ***β***, leading to

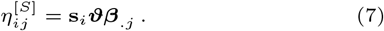

In summary, the coupling of the described features to a shared latent space via the coefficient matrices ***α, δ, ϑ*** allows to quantify to what extent spatiotemporal information influences each taxon’s (latent) abundance after discounting the contribution of the environmental component.

#### Latent modeling of taxon-taxon associations

It is well-established that the abundances of species in an ecosystem are not only driven by environmental or spatiotemporal factors but also by interactions among the species themselves [41]. While discovering detailed ecological interactions among taxa, such as, e.g., competition, mutualism, or commensalism, is beyond the reach of coarse-grained statistical models, VI-MIDAS’ latent space modeling offers a principled mechanism to assess the influence of taxa *co-occurrences* on their respective abundances. We achieve this by borrowing recent ideas from market basket analysis and adopt the so-called SHOPPER utility model for interaction analysis [61]. In SHOPPER, Ruiz et al. [61] proposed a probabilistic model based on the basket data from a supermarket to learn about the latent characteristic of each item and exchangeable/complementary interactions among items. The approach uses item-specific latent variables to define an item-item interaction component. Following their setup, the “interaction”, or, in the biological context, association of the *j*th taxa with any *m*th taxa is given by 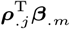 where ***ρ*** = [*ρ*]_*k×q*_ ∈ ℝ^*k×q*^ comprises length-*k* latent variables for each of the *q* taxa. The entries of VI-MIDAS’ interaction component *η*^[*I*]^ for the *j*th taxon in the *i*th sample are thus given by

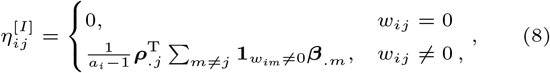

where 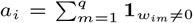 is the total number of taxa present in the *i*th sample. Note that the interaction term ***ρ***^T^***β*** is not symmetric. However, we can derive a symmetrized **I** = [*I*_*i,j*_ ] ∈ ℝ^*q×q*^ with each entry being computed as:

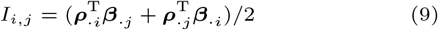

This allows easier downstream network analysis of potentially *positive* (mutualistic) and *negative* (competitive) associations among the taxa, or in our case, among the ecologically relevant clades.

### Variational inference in VI-MIDAS

The generality and flexibility of VI-MIDAS poses a considerable challenge for fast and accurate model parameter estimation. We introduce a variational inference framework that makes estimation in VI-MIDAS feasible and illustrate its performance and parameter sensitivities using the Tara data. For ease of presentation, we summarize the key ingredients below and refer to the extensive Supplementary Information and the documented code base available at https://github.com/amishra-stats/vi-midas) for details.

#### Bayesian model and variational approximation

We begin by denoting all (latent) parameters in the VI-MIDAS framework by ***𝓁*** = {***α, ϑ, β, γ, ρ, τ, Φ***} (see Table S1 of the Supplementary Material). Given the microbial abundance data **W**, the (direct) covariates **X**, and the model parameters **𝓁**, we integrate the generative model (1) into a Bayesian framework where the posterior distribution reads:

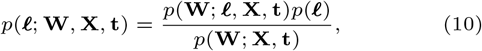

where *p*(**W**; **𝓁, X, t**) = Π_*i,j*_ *p*(*w*_*ij*_ ; *τ*_*j*_ *μ*_*ij*_, *φ*_*j*_) denotes the likelihood of **W** and *p*(**𝓁**) = *p*(***α***)*p*(***δ***)*p*(***β***)*p*(***γ***)*p*(***ρ***)*p*(**Φ**)*p*(***τ***)*p*(***ϑ***) the prior distribution, respectively. To achieve good generalizabilty and interpretability of VI-MIDAS’ over-parameterized model, we place sparsity-inducing Laplace priors with scale parameter *λ* on each of the unconstrained latent variables in the set {***α, δ, β, γ, ρ, ϑ***}. For example, the prior on ***α*** reads *p*(***α***) = Π_*i,j*_ *p*(*α*_*ij*_) with *p*(*α*_*ij*_) = Laplace(0, *λ*). Furthermore, we place an inverse-Cauchy prior on the dispersion parameter **Φ**, i.e., *p*(*φ*_*j*_) = inverse-Cauchy(0, *ν*) and *p*(**Φ**) = Π_*j*_*p*(*φ*_*j*_), and a Uniform(1,2) prior for the shape parameter ***τ***, i.e., *τ*_*j*_ ∼ Beta(1,1) and *p*(***τ***) = Π_*j*_ *p*(*τ*_*j*_). Choosing suitable hyperparameters for the priors will be discussed below.

In the high-dimensional setting, computing the posterior distribution is challenging because of the intractable form of the marginal distribution *p*(**W**; **X, t**) and the non-conjugate priors on the model parameters. Markov Chain Monte Carlo (MCMC) sampling provides a helpful paradigm for obtaining samples from the posterior distribution in the Bayesian framework. However, since MCMC lacks computational efficiency in large/high-dimensional problems, we use mean-field Variational Inference (VI) [34, 72, 8] and approximate the posterior with a variational posterior distribution of the latent variable **𝓁**. Briefly, let *q*(**𝓁 *?***) be the variational posterior distribution with parameter ***ν***. VI approximates sampling of the posterior by minimizing the Kullback-Leibler (KL) divergence,

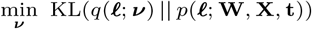

such that supp(*q*(**𝓁; *ν***)) *⊆* supp(*p*(**𝓁; W, X, t**)). It can be shown that the above optimization problem simplifies to maximizing the evidence lower bound (ELBO) given by

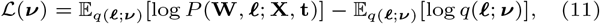

which is a lower bound on the logarithm of the joint probability of the observations log *P* (**W; X, t**) [34]. Replacing the joint distribution *P* (**W, 𝓁; X, t**) with a product of likelihood and prior distribution *P* (**W, 𝓁 X, t**) = *P* (**W**; **𝓁, X, t**)*P* (**𝓁**) further simplifies the objective.

#### Model estimation, hyperparameter tuning, and posterior estimates

The non-convexity of the variational objective and the large number of model parameters require careful assessment of all aspects of model parameter estimation, hyperparameter tuning, and generalization capability. To estimate the parameters of the variational posterior distribution, we employ stochastic gradient descent within the automatic differentiation variational inference (ADVI) framework [36]. The key steps of ADVI are outlined in Algorithm 1 of the Supplementary Material. A prerequisite for model parameter estimation is the identification of suitable model *hyperparameters*. In VI-MIDAS, the key hyperparameters are the scale of the sparsity-inducing Laplace prior, the scale of the inverse-Cauchy prior, and the intrinsic dimensionality *k* of the latent space ***β***, respectively. VI-MIDAS tunes these parameters via random search (see Section 3.3 of the Supplementary Material for details) where the out-of-sample log-likelihood posterior predictive density (LLPD) is used for assessing optimality of the hyperparameters [22]. Due to the non-convexity of the objective and the use of stochastic optimization in VI initialization, we further evaluate the suitability of hyperparameter setting across fifty random initializations and select the hyperparameter set leading to the best averaged LLPD (see Section 3.5 of the Supplementary Material). The computational workflow is implemented in Python using the probabilistic programming language Stan [12] and is available in the GitHub repository (https://github.com/amishra-stats/vi-midas).

After hyperparameter tuning, we re-estimate the final model parameters on complete data. VI-MIDAS generates *m* = 100 posterior samples of each of the latent variables in the set **𝓁** and estimates the model parameters **𝓁** using the mean of the samples from the variational posterior distribution. The model fit is numerically evaluated using the posterior predictive check [60, 22] on the full data. The procedure requires generating *m* posterior samples, denoted by the random variables 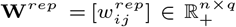, and then computing the p-value of the model fit as p-value := *p*(*t*(**W**^*rep*^) *< t*(**W**)), where *t* is the test statistic. In practice, we use the test statistics *t*(**W**^*rep*^) = **E**(log *p*(**W**^*rep*^|**𝓁**)) and *t*(**W**) = **E**(log *p*(**W**|**𝓁**)).

## Results

### VI-MIDAS recapitulates broad statistical patterns of the observed species abundances

VI-MIDAS’ hyperparameter tuning revealed that the setting *k* = 200, *λ* = 0.246, and *ν* = 0.10063 achieved the highest average LLPD of 3.332 on the Tara data (see Figure S7 in the Supplementary Material). For this setting, a posterior predictive check on the generated samples achieved a p-value = 0.53. We thus fail to reject the null hypothesis that the posterior samples are different from the observed **W**. Figure 3a and 3b the observed and estimated abundance profiles (averaged over *m* = 100 samples), respectively. Figure 3c shows the count histograms of data and model (pooled across all samples and species), and Figure 3d the Q-Q plot. We observe that, apart from the low-abundance tail of the distribution, VI-MIDAS broadly recapitulates the statistical abundances patterns across all samples and species.

**Fig. 3.**
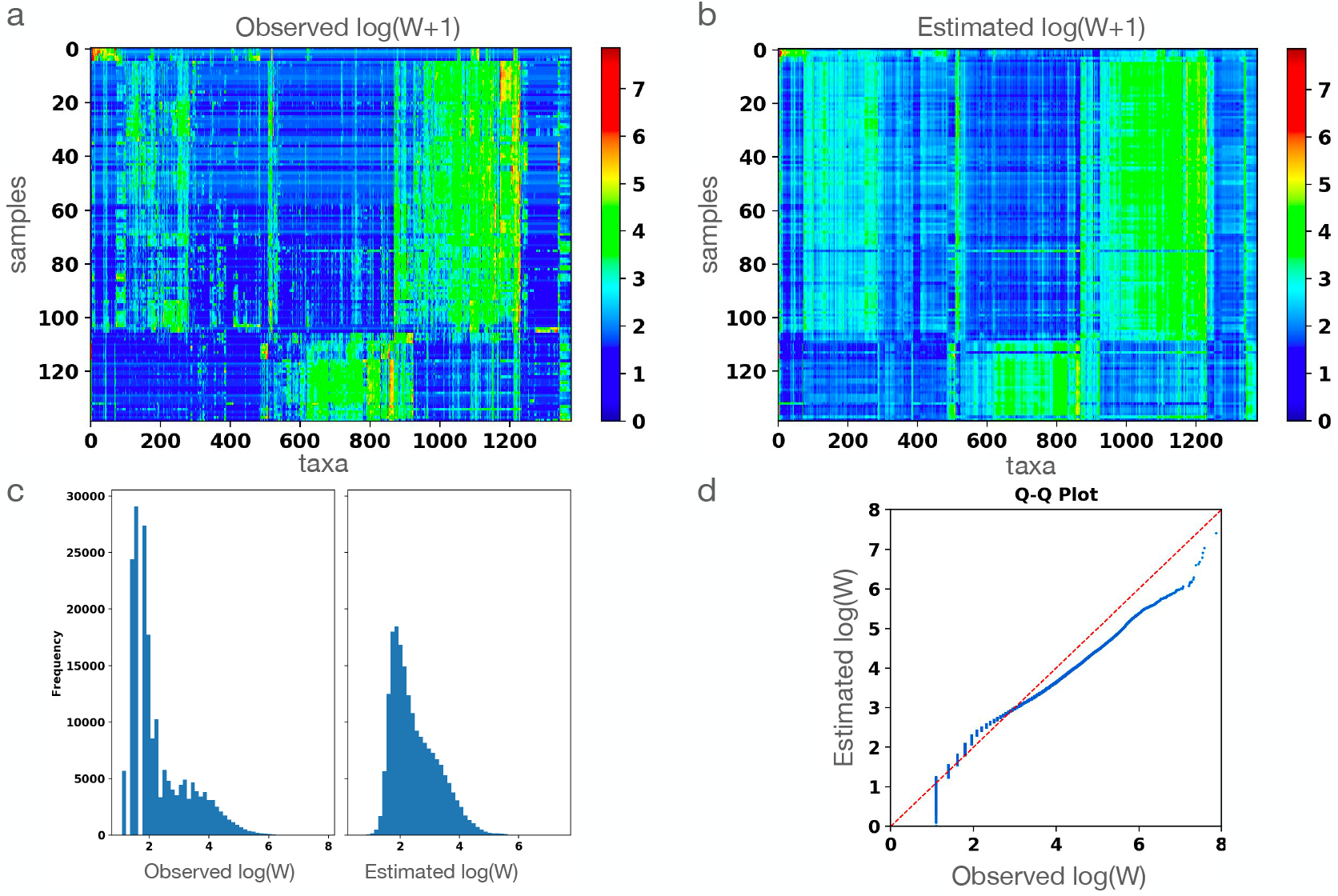
Comparison of observed abundances and VI-MIDAS posterior samples: **a**. Heatmap showing the abundance profile log(**W** + 1) of 1378 species for *n* = 139 samples. **b**. Expected value of the abundance using the hyperparameter corresponding to best model fit. **c**. Histograms of observed and estimated species abundances. **d**. Q-Q plot comparing the observed and estimated abundance profile of the species.

### VI-MIDAS identifies depth and environmental features as main drivers

We next assessed the contribution of each model component toward explaining the species abundance patterns in the Tara data. The modularity of the VI-MIDAS framework facilitates an “ablation” study (see Section 3.4 of the Supplementary Material) where each model component is excluded, followed by a re-evaluation of the out-of-sample LLPD. Table S4 (see Supplementary Materials) shows the LLPDs of the full model and the model after ablation of the environmental(E), province(P), ocean depth(D), seasonality(S), and latent interaction (I) component, respectively.

Firstly, the ablation study confirmed that all components helped improve model generalization since every ablated model has reduced out-of-sample LLPDs. While the seasonality component(S) shows comparatively little influence on explaining the abundance pattern in the current model, as previously observed for this dataset [67], the out-of-sample LLPD is reduced the most when the ocean depth(D) component is ablated (LLPD=-3.3882). This reflects the well-known depth stratification of marine species between the sunlit ocean and aphotic deep ocean ecosystems. Figure S3 in the Supplementary material illustrates the learned depth stratification across all taxa, as reflected in the component *δ****β***. The environmental component was identified as the second most important component with an LLPD reduction of -3.3554.

Figure 4 summarizes the estimated effects ***δβ*** of the ocean depth features and the environmental effects ***γ*** on the abundance of species aggregated into ERCs, respectively. The ocean depth summary (Fig. 4a) reveals three distinct sets of occurrence patterns for two different groups of ERCs.

**Fig. 4.**
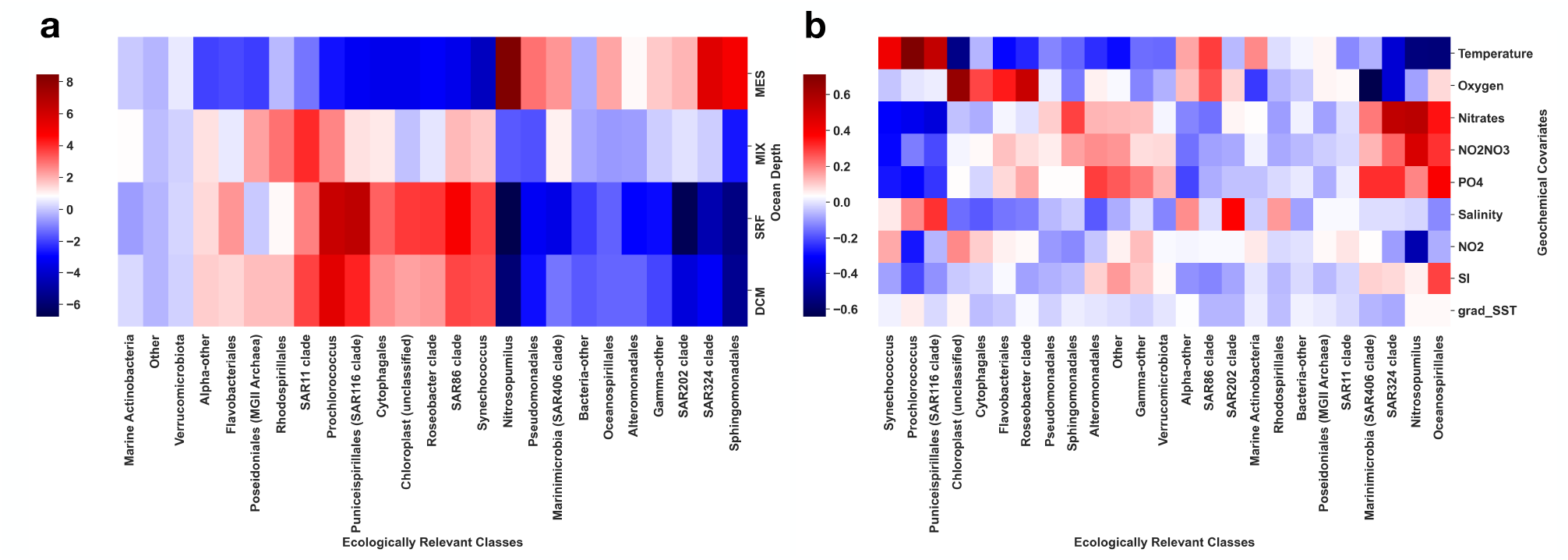
Summary of the estimated average effect sizes of the influence of **a**. ocean depth (VI-MIDAS model component *δ****β***) and **b**. environmental covariates (VI-MIDAS model component *γ*) on all ecologically relevant classes (ERCs).

One group (right most in Fig. 4a) comprises ERCs such as Nitrosopumilius, Pseudomonadales, SAR 324 clade, and Sphingomonadales which thrive in the Mesopelagic (MES) zone. A second group includes species like Prochlorococcus, SAR 116 clade, and Synechococcus, which flourish within the ecosystem of the ocean’s Deep Chlorophyll Maximum (DCM) and Surface Mixed Layer (SRF) zones. The third group comprises marine Actinobacteria, Verrucomicrobiota, and others that show no dependence on depth. A summary of geochemical features highlights temperature (the top row in Fig. 4b) as the primary positive factor influencing the abundance of Synechococcus, Prochlorococcus, and Puniceispirillales (SAR116 clade). Oxygen concentration emerges as the main positive driver of abundance for Cytophagales, Flavobacteriales, and Roseobacter clades, while Nitrates, Nitrites, and Phosphate are identified as key drivers for the SAR324 clades, Nitrosopumilus, and Oceanospirillales (four right most columns in Fig. 4b). The estimated patterns broadly recapitulate known biology about ocean microbial ecosystems.

### VI-MIDAS reveals five latent microbial sub-communities

The generative model (1) of VI-MIDAS includes the taxon-specific latent variables ***β*** ∈ ℝ^*k×q*^ to integrate spatiotemporal features and taxon-taxon associations. For the Tara data, VI-MIDAS’ hyperparameter tuning scheme identified *k* = 200 as best latent dimension. After model estimation, the resulting *k−*dimensional latent vectors can be thought of as representing the hidden *marginal* characteristics of each of the *q* taxa after discounting spatiotemporal and species-species association effects, and adjusted for environmental covariates. The latent space representation thus provides an excellent opportunity to partition the different taxa into coherent sub-groups (or modules) that likely reflect functionality or niche occupation in the global ocean, independent of environmental, taxonomic or phylogenetic relatedness.

To quantify similarity between microbial taxa in the latent space, we first computed cosine distances of all pairs of the *q* latent vectors. This particular choice of distance allows us to bypass the non-identifiability issue of the parameter ***β***. We used the resulting distance matrix to construct a k-nearest neighbors graph (*k*_nn_ = 10). Figure 5 shows the latent space embedding using a force-directed layout of the k-nn graph. The latent space representation reveals several distinct microbial sub-communities, dominated by a few ERCs, including one sub-community dominated by Prochlorococcus and SAR11 clades and one dominated by Nitrosopumilus. We next performed Clauset-Newman-Moore greedy modularity analysis of the nearest neighbor graph [15] and identified five distinct modules in the latent space (see M1-M5 in Fig. 5 with top five ERCs highlighted and color-coded). Module 1 (M1) comprises Flaviobacteriales, SAR86 clades, and the Chloroplast class. SAR11 clade, SAR86 clade, and Flavobacteriales are heterotrophs with functional similarity in oxidizing carbon in the ocean [3]. Both SAR86 clade and SAR11 clade follow a similar seasonal pattern (in the Bermuda Atlantic Time Series oceanographic stations) and coexist in oligotrophic regions with less nutrient supply [73]. Module 2 (M2) includes Nitrosopumilus, Marinimicrobia, and SAR324 clades. Existing literature supports that SAR11 clade (a subgroup of a species), Marinimicrobia, and MGII Archaea are more abundant in deep sea water [76]. Module 3 (M3) comprises Prochlorococcus, SAR11, Marine Actinobacteria, and SAR86 clades, among others, all comprising dominant taxa of the sunlit ocean. The two smallest modules 4 and 5 (M4 and M5, respectively) are dominated by Alteromonadales and are separating M2 from M1 and M3. Interestingly, Module 4 also comprises Synechococcus species. This module thus hints at the known metabolic dependency of certain Alteromonadales taxa on Synechococcus (a photoautotroph) [80]. Although the latent representation does separate the majority of ERCs into distinct subgroups, we nonetheless observe that taxa of certain ERCs are spread out over the latent space, indicating different niche specialization. For instance, the SAR11 clade, one of the most abundant marine microbial taxa, is present in three different modules. Likewise, taxa in the SAR86 clade are present in both modules M1 and M3. For ease of identification, Table S3 summarizes each module in terms of the composition of the ERCs and their abundance.

**Fig. 5.**
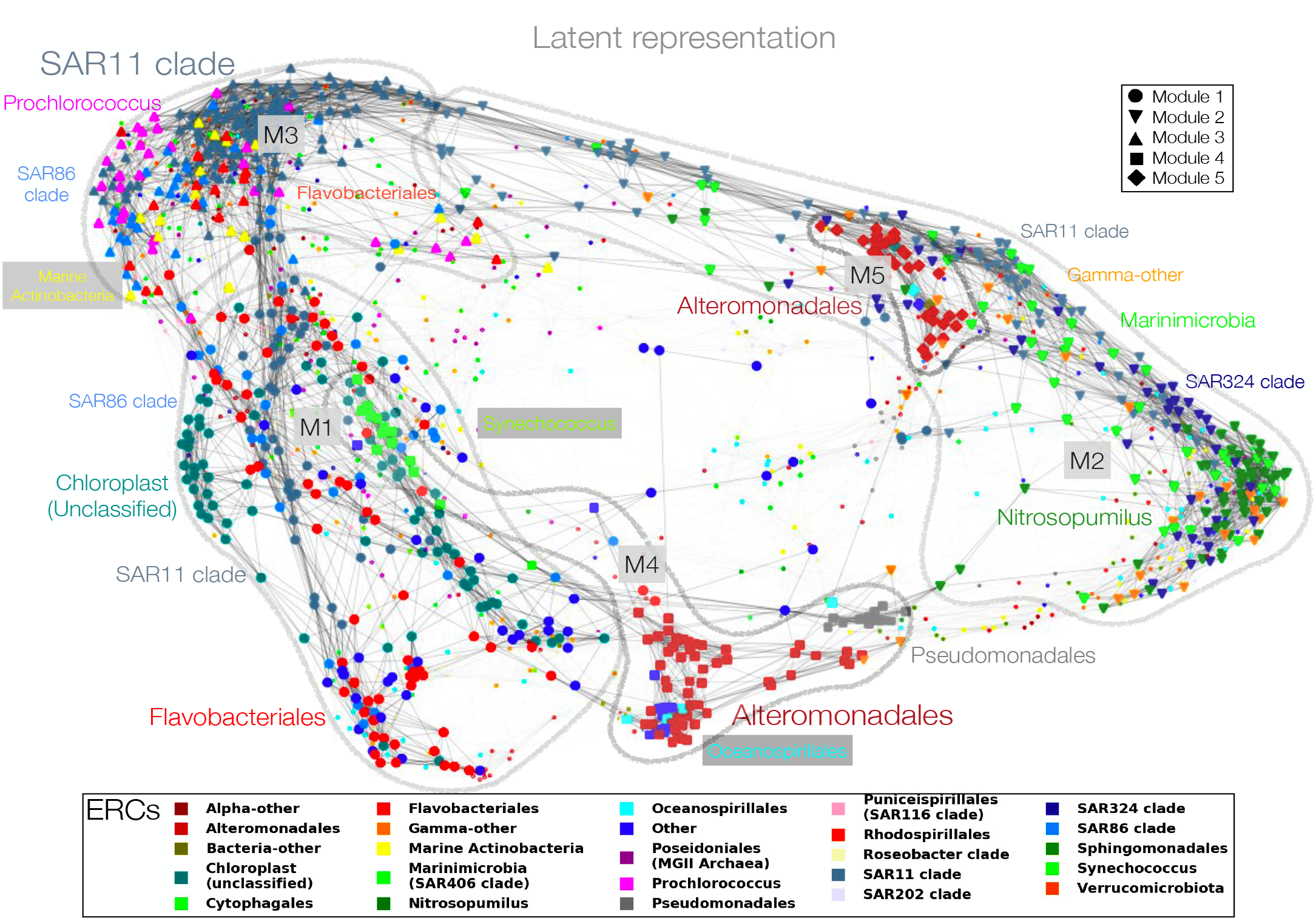
Low-dimensional embedding of the latent representation ***β*** using a k-nearest-neighbor (*k*_nn_ = 10) graph of cosine distances. Modularity analysis reveals five distinct graph modules. We highlight 825 out of a total of 1378 taxa, comprising the top five ERCs (color-coded) in each of the five modules (see main text for further information).

### Global associations between biogeography and latent microbial sub-communities

VI-MIDAS’ integrative model also enables a quantitative description of the identified microbial sub-communities in terms of the direct and indirect coupling covariates. Figure 6 illustrates how the compositions of ERCs in each of the five modules are related to the most important environmental and spatial covariates.

**Fig. 6.**
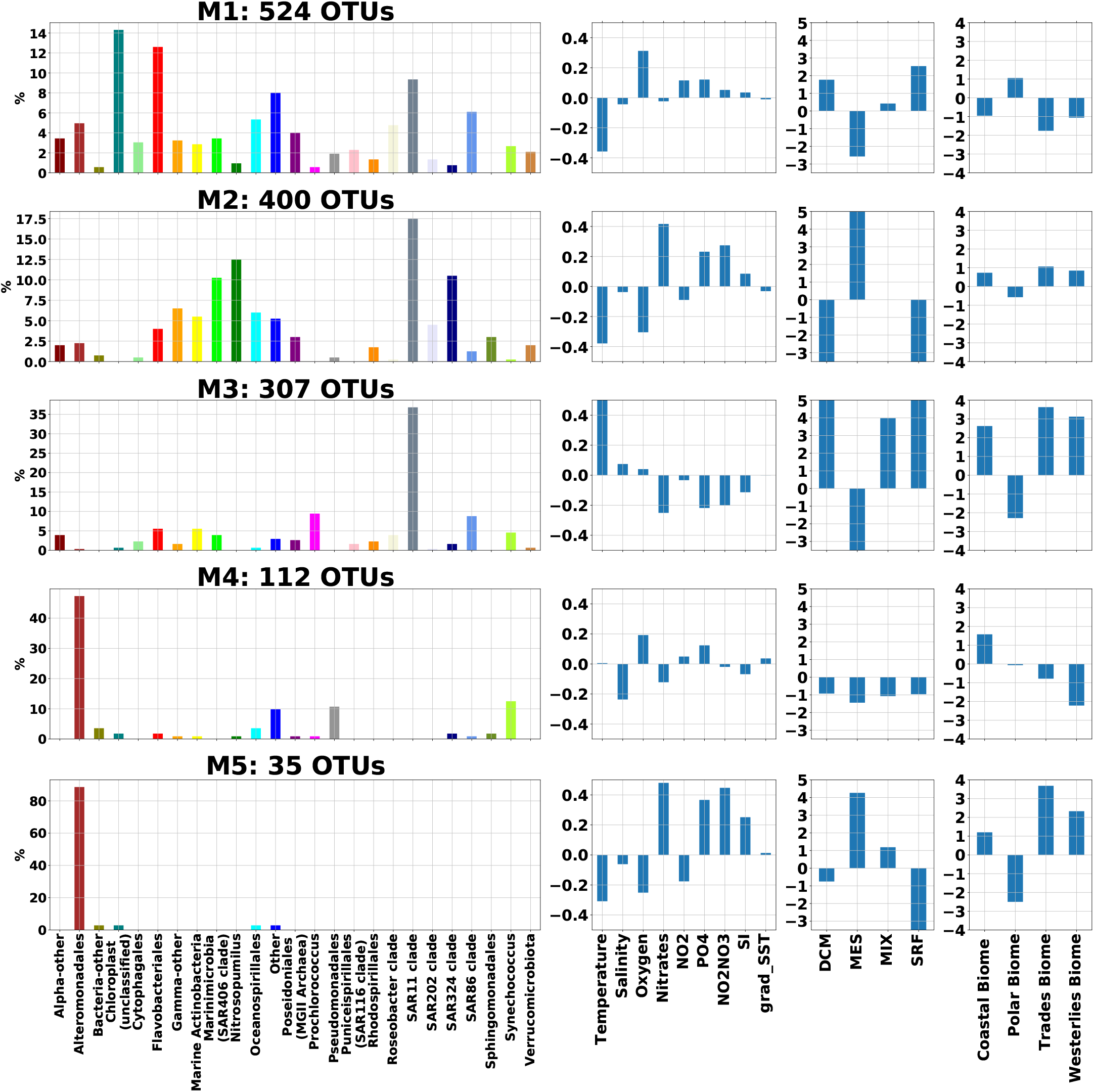
Global associations between biogeography and covariates: Each row presents the average effect size of the association between the microbial abundances of taxa in a module (M1-M5) to the geochemical features, ocean depth and province/location (from left to right). A module (leftmost) is shown as the composition (in %) of the ERCs. Each module comprises different number of taxa *{*524, 400, 307, 112, 35*}*, respectively. Modules M1-M3 cover the majority of taxa, and M4-M5 two smaller Alteromonadales-dominated sub-communities.

Using the mean of the posterior sample from the VI-MIDAS model, we used the estimated ***γ*** as the effect sizes of the environmental features **X, *δβ*** as effect sizes of depth, and ***αβ*** as the effect sizes of the *r* provinces, respectively Figure 6 reports the average effect sizes of association to the four modules.

The module M1 represents taxa coexisting in the SRF and DCM zone of the ocean. The abundance of taxa in the module is associated with a higher concentration of oxygen, PO_4_, and NO_2_NO_3_ and lower temperature and salinity. In addition to representing the taxa SAR11 clade, SAR86 clade, Chloroplast, and Flavobacteriales, the module also includes Synechococcus, Oceanospirillales, and Poseidoniales. Synechococcus is a unicellular prokaryotic autotrophic picoplankton that participates in the marine ecosystem as a primary producer via photosynthesis. Similarly, Chloroplast sequences are a signature of eukaryotic phytoplankton, though their host eukaryote is not identified in the TARA Oceans dataset. The presence of both taxa in M1 thus is consistent with environments that have higher oxygen concentrations due to photosynthesis and gas exchange with the atmosphere.

Module M2 mainly represents the species coexisting in the MES zone (200 m to 1000 m) of the ocean (see Figure 2 (e)). M2 almost exclusively represents the ERCs Nitrosopumilus and SAR324 clade. The abundance of the species in the group is associated with a lower concentration of oxygen and temperature, and higher concentrations of nitrates, PO_4_, and NO_2_NO_3_. In the oxygen-depleted environment, Nitrosopumilus survives by oxidizing ammonia to nitrite, confirming the observed association pattern [6]. Marinomicrobia (SAR406 clade) in groups M1 and M2 allow us to distinguish subgroups of species that can survive in both deep and shallow water [76].

Module M3 comprises the highest mean abundance of all taxa is highest, primarily representing the taxa SAR11 clade, SAR86 clade, and Prochlorococcus (cyanobacteria). The abundance of the species in the group is positively associated with depth indicators {DCM, MIX, SRF} and negatively associated with MES. Among the geochemical factors, temperature, salinity, and oxygen concentration are positively associated, whereas the concentration of nitrates, PO_4_, and NO_2_NO_3_ is negatively associated with the taxa.

Module M4 primarily represents Alteromonadales (Proteobacte and some Pseudomonadales (Proteobacteria) and Synechococcus. Their abundance is associated with factors such as lower salinity and higher oxygen concentration. Module M5 also primarily represents Alteromonadales. Based on its association with the ocean depth indicators and geochemical features, we conclude that these taxa can survive in a deep-sea environment characterized by lower temperatures and oxygen concentrations. Associative patterns of Alteromonadales in M4 and M5 differ significantly, suggesting distinct ERC sub-groups that populate different niches.

### Positive and Negative interactions among ERCs

VI-MIDAS includes a mechanism for learning microbial interactions adjusted for direct (here, environmental) covariates. Contrary to prominent (partial) correlation-based methods [21, 38], VI-MIDAS follows the SHOPPER utility model [61] and quantifies pairwise interactions **I**_*ij*_ between any two taxa *i* and *j* in terms of the latent variables ***ρ*** and ***β*** (see Eq. 9).

To get a high-level view of the estimated interactions, we aggregated the adjacency matrices of significant positive and negative interactions among taxa by ERCs (for a more detailed view of the most significant taxon-level interactions, we refer to Section 4 of the Supplementary Materials). Figure 7 illustrates the aggregated positive (lower triangle) and negative (upper triangle) interactions among ERCs. The diagonal entry highlights the maximum of the two types of interactions to avoid confusion (see also Section 4 of the Supplementary Materials for the matrix of ratios between positive and negative interactions). We observe that SAR11 clade and Rhodospirillales form positive interactions with almost all other ERCs. SAR11 clade and Rhodospirillales belong to the Alphaproteobacteria phylum that play a critical role in carbon and nitrogen fixation [40, 51], potentially explaining the large number of interactions. However, members of the SAR11 clade also form many negative interactions with other ERCs.

**Fig. 7.**
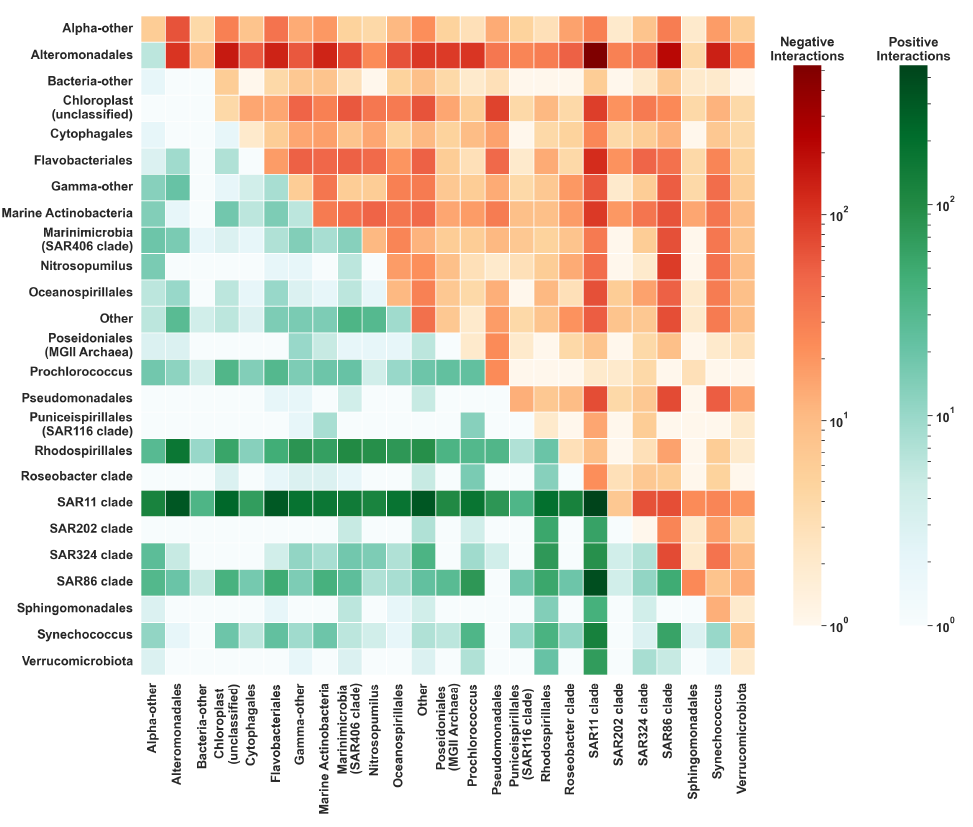
Summary of taxonomic interactions: The adjacency matrices of significant positive and negative interactions among taxa are grouped and aggregated by their ERCs type. Interactions summary by the ERCs types. Lower triangle reports positive interactions, the upper triangle reports negative interactions. Diagonal entries show the maximum of either (positive or negative) self-interaction.

Alteromonadales exhibits primarily negative interactions with other ERCs (the strongest one with SAR11).

## Discussion

In recent years, multimodal and multi-omics microbiome survey data have emerged for a wide range of microbial habitats [68, 67, 47, 27, 65, 1]. These data collections hold the promise to describe and understand the functional interplay between the underlying microbial ecology and the host or the environment the microbiota resides in. Learning interactions among species and habitat characteristics from observational data remains, however, a challenging problem. To this end, we have proposed VI-MIDAS (Variational Inference for MIcrobiome survey DAta analysiS), a flexible and efficient probabilistic framework for microbiome survey data analysis.

VI-MIDAS uses the negative binomial distributional framework in combination with a principled centering transformation to model overdispersed amplicon abundance data and comprises three mechanisms to integrate concomitant covariate data into the generative model: (i) a direct coupling mechanism, (ii) an indirect latent coupling mechanism, and (iii) a latent interaction term. These terms are linearly linked to the probability distribution’s mean parameter. Because of the intractable form of the marginal distribution of data, we apply mean-field variational inference framework to learn an approximate posterior distribution of the parameters.

VI-MIDAS is available in Python and uses the probabilistic programming language Stan [12]. The implementation is available on GitHub (https://github.com/amishra-stats/vi-midas). The repository also includes Python scripts and Jupyter notebooks for VI-MIDAS’ three-stage parameter estimation framework: hyperparameter tuning, component contribution analysis, and sensitivity analysis.

To illustrate the VI-MIDAS modeling and analysis workflow, we have used data from the global Tara expedition [67], connecting the available spatiotemporal and environmental characteristics with generative modeling of the amplicon count data. To ease interpretability, we also grouped the amplicon-derived taxa into expert-annotated ecologically relevant classes (ERCs) which may be of independent interest for the analysis of other marine sequencing data. Focusing on the *q* = 1378 most abundant taxa representing 23 ERCS, we integrated the geochemical data using the direct coupling mechanism, effectively removing influence of common environmental factors such as temperature, salinity, and elemental compositions on microbial abundances. The remaining spatiotemporal features, including season, ocean province, and depth, as well as species-species associations are integrated through the latent coupling and interaction mechanism, thus delivering a latent species representation, adjusted for the influence of all available covariates. The learned VI-MIDAS’ model thus not only provides a convincing generative count model for the Tara data but also allows integrated statistical analysis of covariate feature effects and taxa abundances.

Modularity analysis of the similarity network of VI-MIDAS’ latent species representation revealed that the majority of taxa (¿1200) can be categorized into three global microbial communities (M1-M3 in Figure 5), including a low-temperature/high-oxygen community (M1), dominated by Flavobacteriales and the Chloroplast ERC, a mesopelagic community (M2) dominated by SAR11, SAR324, and Nitrosopumilus, and a high-temperature community (M3) dominated by SAR11 and Prochlorococcus, the later of which is the most abundant clade in the oligotrophic subtropical and tropical oceans (see e.g., [66] and references therein). Furthermore, our analysis suggests two distinct Alteromonadales-dominated communities that show different depth and province dependencies (M4-M5) (see Figure 6 for further global associations overview). It is noteworthy that Alteromonadales also play a pivotal role in the latent interaction analysis, showing widespread negative associations with other ERCs. We posit that the potentially distinct role of Alteromonadales in the global ocean might be of interest for follow-up analysis on other data sets, including recent data on the global mesopelagic zone [59].

While our ablation study showed evidence that all VI-MIDAS components for the Tara data contribute to the quality of the generative model, the model is just one of several available alternatives. For covariate inclusion, we deliberately chose to directly adjust the microbial abundances for geochemical covariates to better carve out “hidden” relationships among the species. Nonetheless, the VI-MIDAS framework naturally enables other model constructions. For instance, one could have removed the direct coupling component and link all concomitant features to the latent space representation, or alternatively, remove the latent representation altogether and directly adjust for all covariates. We will explore such modifications in future studies. Moreover, while we chose the Negative Binomial model as base distribution for the most abundant taxa, the variational formulation lends itself to other statistical models for microbial count data, including zero-inflated or hurdle-type extensions of the Negative Binomial model [19] or the Dirichlet-Multinomial model [30, 53]. Finally, in its current state, VI-MIDAS is built on Stan [12] with tailored Python code for optimization, model selection, and analysis. The advent of extensive statistical packages in modern deep learning tools, such as Tensorflow distributions [17] or PyTorch [55], may enable efficient porting of VI-MIDAS into these general-purpose ecosystems. Paired with variational inference tools [35], would potentially allow for faster model adaptation and alternative optimization routines. In summary, VI-MIDAS provides a novel probabilistic framework for learning environment- or host-specific feature associations, latent species characterization, and species-species interactions from microbiome survey data. With minimal adjustment, the framework is readily available for the analysis of other large-scale survey data, including gut microbiome surveys [33, 45, 20], thus representing a potentially valuable general-purpose tool for the integrated analysis of modern microbiome data collections.

## Data availability

We have used microbial species abundance data from the Tara Ocean Expedition, available at (http://ocean-microbiome.embl.de/companion.html).

## Code availability

The source code required to reproduce the results in this article is freely available at (https://github.com/amishra-stats/vi-midas).

## Footnotes

1 http://ocean-microbiome.embl.de/companion.html

