## Supplement figures for "Variational inference for microbiome survey data with application to global ocean data"

March 18, 2024

#### 1 Supplementary Figures

We report the supplementary figures of the data analysis related to the main manuscript.

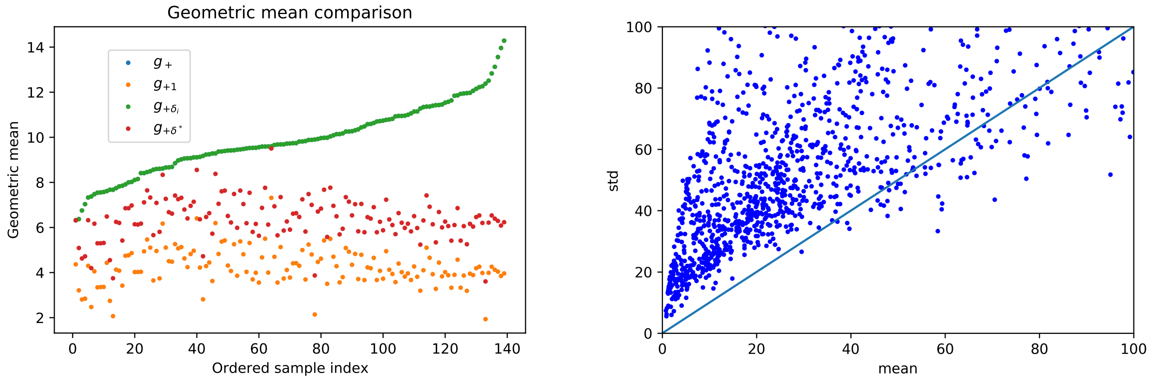


(a) (b)

Figure S1: a) Compares the possible approach of computing the geometric mean of the microbial abundance sample to be considered as an offset term $\mathbf{t}$ in the VI-MIDAS. Suggested approach mainly differs in the pseudo $\delta_i$ added to zero entries in a given sample; b) Over-dispersion of abundance is demonstrated by comparing the means and standard deviations of the abundance of $q = 1378$ species selected for analysis. According to the analysis, the negative binomial distribution is an appropriate distribution to represent the microbial abundance data.

*Corresponding author

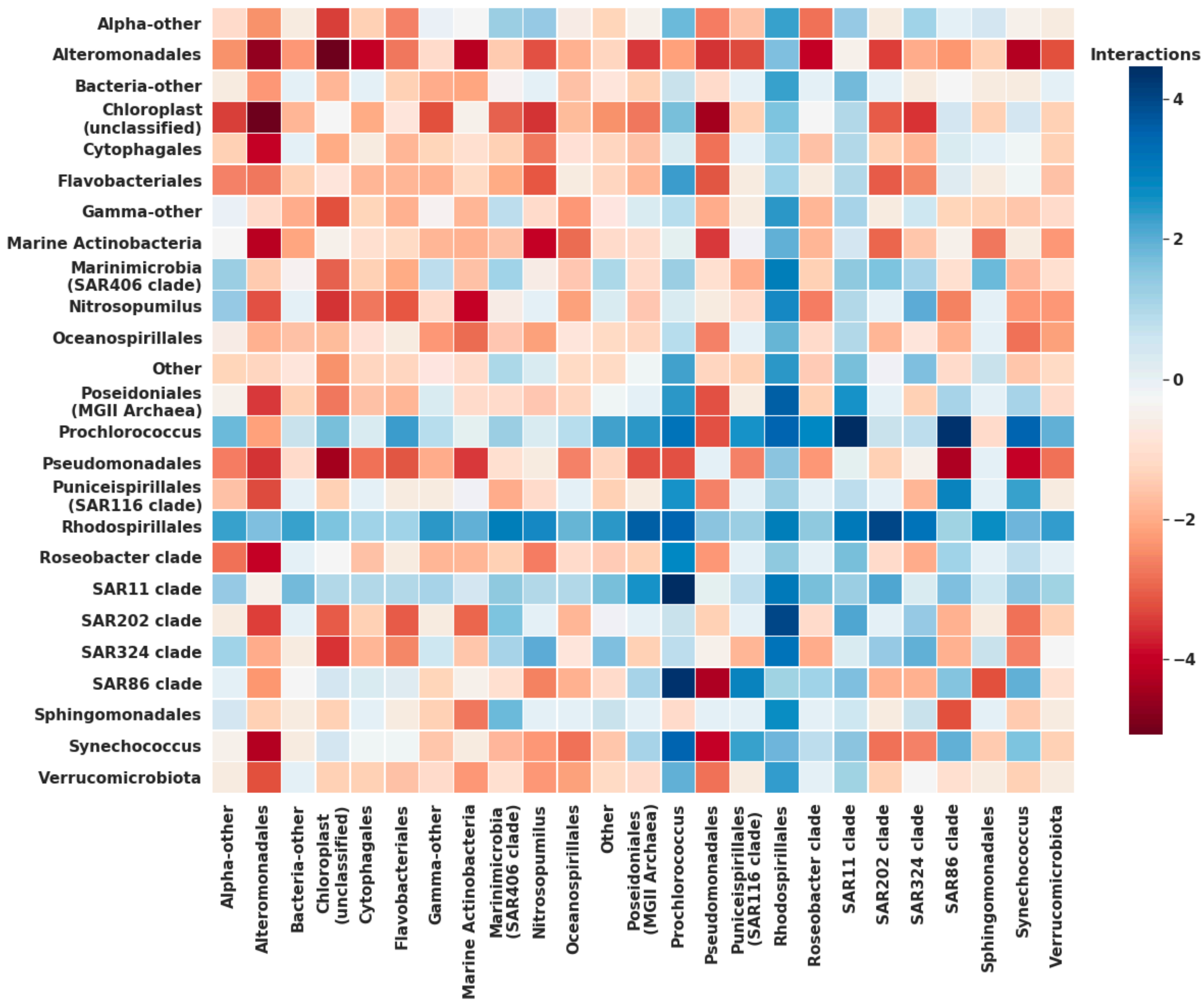


Figure S2: Comparison of interactions among ERC identifiers: The adjacency matrices of significant mutualistic and competitive interactions among OTUs are grouped and aggregated by their ERC identifier. A block represents the logarithmic ratio of the number of mutualistic interactions versus the number of competitive interactions. Positive and negative values, accordingly, represent one larger than the other. Blue for mutualistic interaction and Red for competitive interactions.

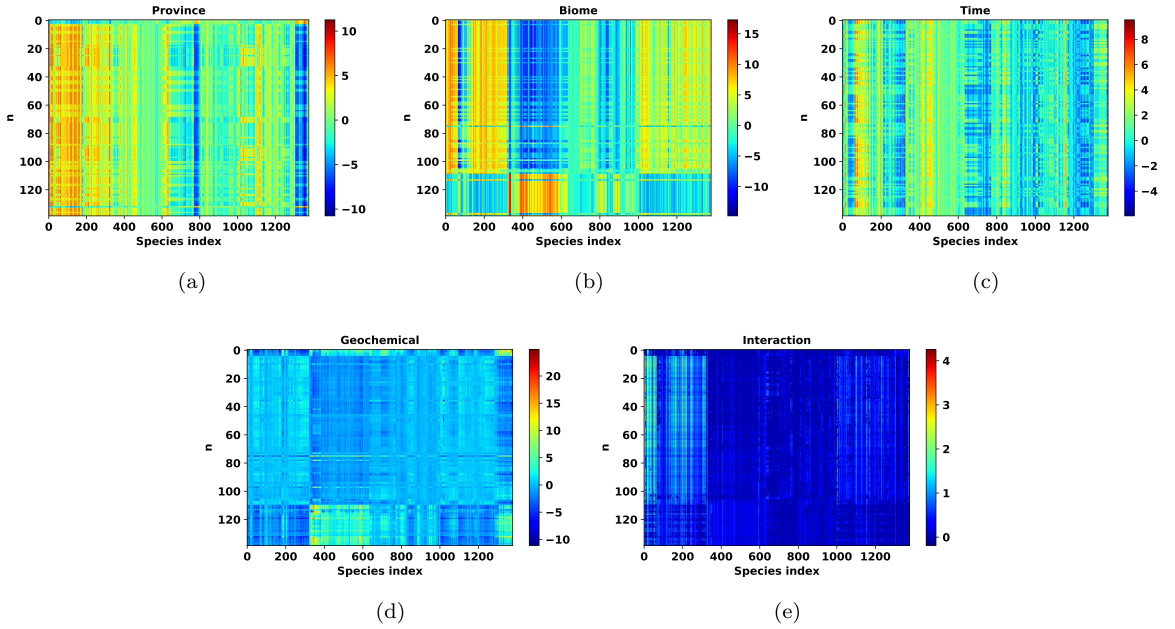


Figure S3: Model fit: Estimate of the non-interaction components in the linear predictor $\boldsymbol{\eta}$: a) Province indicator; b) Biome (depth) indicator; a) Quarter (time) indicator; d) Biogeochemical factor; e) Embedding effect.

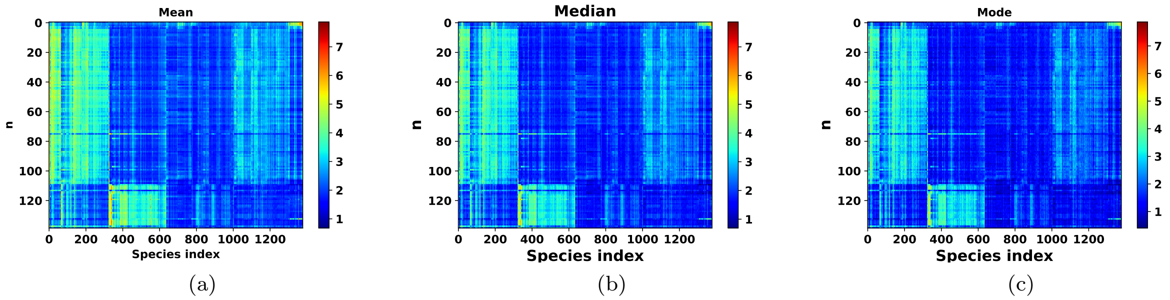


Figure S4: Model fit: a) Mean estimate of the posterior sample using VI-MIDAS; b) Median estimate of the posterior sample using VI-MIDAS; a) Mode estimate of the posterior sample using VI-MIDAS.

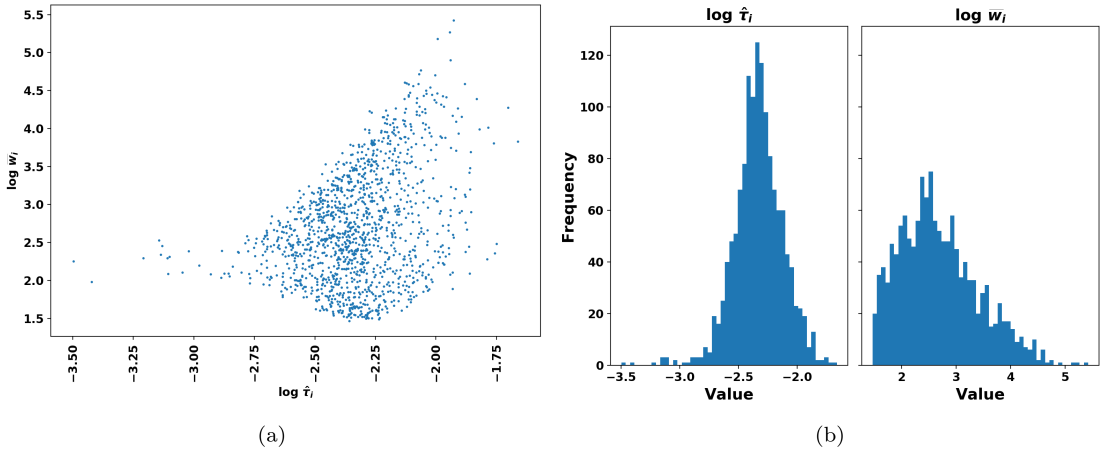


Figure S5: a) Scatter plot comparing entries of the CLR transform of the observed and estimated microbial species abundance given by $\log \mathbf{W}/\mathbf{T}$ and $\log \mathrm{E}(\mathbf{W})/\mathbf{T}$, respectively; b) Comparison of the estimate of $\tau$ and mean of the species abundance.

### 2 Supplementary Tables

Table S1: Dimension of the latent variables in the parameter set $\boldsymbol{\ell}$ of the VI-MIDAS model.

| | $\boldsymbol{\gamma}$ | $\boldsymbol{\beta}$ | $\boldsymbol{\rho}$ | $\boldsymbol{\alpha}$ | $\boldsymbol{\delta}$ | $\boldsymbol{\vartheta}$ | $\boldsymbol{\tau}$ | $\boldsymbol{\Phi}$ |
|---|---|---|---|---|---|---|---|---|
| # ncols | 1378 | 1378 | 1378 | 4 | 4 | 4 | 1378 | 1378 |
| # nrows | 11 | 200 | 200 | 200 | 200 | 200 | 1 | 1 |

Table S2: Selected value of the latent variable dimension $k$ and the hyper-parameters $\{\lambda, \upsilon\}$ of the Laplace and Inverse-Cauchy priors in the range of one standard deviation of the largest value of the $\mathrm{LLPD}_o$.

| Index | k | $\lambda$ | $\upsilon$ | $\mathrm{LLPD}_o$ |
|---|---|---|---|---|
| 1 | 200 | 0.246 | 0.10063 | -3.318726 |

Table S3: Top five ecologically relevant classifications (ERC) indicator of taxa in the five modules identified in the network (shown in Figure ??) highlighting similar nodes. The table reports the composition (as %) of 825 taxa in terms of the ERC indicator and their mean abundance.

| Module | ERC | Abundance (Mean) | Members (%) | Module | ERC | Abundance (Mean) | Members (%) |
|---|---|---|---|---|---|---|---|
| | Chloroplast | 12.49 | 9.09 | | Alteromonadales | 7.24 | 6.42 |
| | Flavobacteriales | 11.00 | 8.00 | | Synechococcus | 27.70 | 1.70 |
| 1 | SAR11 clade | 19.44 | 5.94 | 4 | Pseudomonadales | 4.83 | 1.45 |
| | Other | 10.77 | 5.09 | | Other | 6.37 | 1.33 |
| | SAR86 clade | 18.91 | 3.88 | | Oceanospirillales | 5.78 | 0.48 |
| | SAR11 clade | 18.67 | 8.48 | | Alteromonadales | 17.84 | 3.76 |
| | Nitrosopumilus | 23.03 | 6.06 | | Bacteria-other | 12.22 | 0.12 |
| 2 | SAR324 clade | 15.48 | 5.09 | 5 | Chloroplast | 17.81 | 0.12 |
| | Marinimicrobia | 16.69 | 4.97 | | Oceanospirillales | 36.30 | 0.12 |
| | Gamma-other | 11.13 | 3.15 | | Other | 13.29 | 0.12 |
| | SAR11 clade | 45.63 | 13.70 | | | | |
| | Prochlorococcus | 53.06 | 3.52 | | | | |
| 3 | SAR86 clade | 34.74 | 3.27 | | | | |
| | Flavobacteriales | 28.33 | 2.06 | | | | |
| | Marine Actinobacteria | 29.17 | 2.06 | | | | |

### 3 Methods Details

#### 3.1 Controlling for the relative abundance data in VI-MIDAS

Due to experimental limitations and systematic bias in high-throughput sequencing, we do not have access to microbial species' absolute (actual) abundance data. Several analysis techniques have been proposed for use on the transformed data. Some acceptable transformation techniques include centered log-ratio transform (CLR), relative abundance, and isometric log-ratio transform [Callahan et al., 2016]. Many statistical methods [Shi et al., 2016, Mishra et al., 2019] consider the relative abundance (compositional) data for exploratory and predictive tasks in microbiome data analysis. For example, CLR transformation leads to centering the log of abundance by its geometric mean, i.e.,

$$\mathrm{CLR}(\mathbf{w}) = \log \mathbf{w} - \log g(\mathbf{w})$$

where $g(\mathbf{w})$ is the geometric mean of $\mathbf{w}$. Instead of using the relative abundance data for a predictive model, VI-MIDAS overcomes the limitations of the microbial abundance data by the use of a suitable offset term. By including $t_i = \log g(\mathbf{w}_i)$ as an offset term in the log-link function (2) of VI-MIDAS, one can control for such limitations; see [Zhang et al., 2017].

A significant number of entries in the abundance matrix $\mathbf{W}$ are zeros. Some of these may reflect the absence of species, while others may be caused by experimental limitations. Hence, one may consider computing the geometric mean using only non-zero entries in a sample ($g_+$) or after adding a pseudocount ($\delta_i$) to the $i$th sample. A standard approach is to use $\delta_i = 1$, denoted $g_{+1}$. In the VI-MIDAS model, we mainly follow de la Cruz and Kreft [2018] to compute the pseudocount $\delta_i$ by solving

$$\delta_i = \sup\{\delta^* \in (0,\infty) \mid G_{\mathbf{w}_i,\epsilon}(\mathbf{w}_i) - g(\mathbf{w}_i^+) \leq \epsilon g(\mathbf{w}_i^+)\},$$

where $G_{\mathbf{w}_i,\epsilon}(\mathbf{w}_i) = \exp\left(\frac{1}{n}\sum_{j=1}^{q} \log(w_{ij} + \delta_i)\right) - \delta_i$, $\mathbf{w}_i^+ = \{w_{ij} | w_{ij} > 0\}$ is the set of nonzero entries in $\mathbf{w}_i$ and $\epsilon$ is the relative difference between the standard geometric mean $g(\mathbf{w}_i^+)$ and our modified geometric mean. Let us denote the geometric mean by $g_{+\delta_i}$. In practice, one may also use $\delta^* = \min_{i=1}^{n} \delta_i$ as a possible alternatives with geometric mean as $g_{+\delta^*}$. Figure S1 (a) in the supplementary materials compares the geometric mean of $n$ samples using the suggested approach.

##### 3.2 Variational inference for estimation

VI-MIDAS is an over-parameterized model. To generalize well on the test data, VI-MIDAS uniformly places a Laplace prior with scale parameter $\lambda$ on each of the unconstrained latent variables in the set $\{\boldsymbol{\alpha}, \boldsymbol{\delta}, \boldsymbol{\beta}, \boldsymbol{\gamma}, \boldsymbol{\rho}, \boldsymbol{\vartheta}\}$, i.e., $p(\alpha_{ij}) = \text{Laplace}(0, \lambda)$ and $p(\boldsymbol{\alpha}) = \prod_{i,j} p(\alpha_{ij})$. Also, we place a inverse-Cauchy priors on the dispersion parameter $\boldsymbol{\Phi}$, i.e., $p(\phi_j) = \text{inverse-Cauchy}(0, \upsilon)$ and $p(\boldsymbol{\Phi}) = \prod_j p(\phi_j)$, and a Uniform(1,2) prior for the shape parameter $\boldsymbol{\tau}$, i.e., $\tau_j \sim \text{Beta}(1,1)$ and $p(\boldsymbol{\tau}) = \prod_j p(\tau_j)$. Given the microbial abundance data $\mathbf{W}$, the geochemical covariates $\mathbf{X}$, the model parameter $\boldsymbol{\ell}$ and the generative model (1), we express the posterior as

$$p(\boldsymbol{\ell}; \mathbf{W}, \mathbf{X}, \mathbf{t}) = \frac{p(\mathbf{W}; \boldsymbol{\ell}, \mathbf{X}, \mathbf{t}) p(\boldsymbol{\ell})}{p(\mathbf{W}; \mathbf{X}, \mathbf{t})}, \tag{1}$$

where $p(\mathbf{W}; \boldsymbol{\ell}, \mathbf{X}, \mathbf{t}) = \prod_{i,j} p(w_{ij}; \tau_j \mu_{ij}, \phi_j)$ is the likelihood of $\mathbf{W}$ and $p(\boldsymbol{\ell}) = p(\boldsymbol{\alpha})p(\boldsymbol{\delta})p(\boldsymbol{\beta})p(\boldsymbol{\gamma})p(\boldsymbol{\rho})p(\boldsymbol{\Phi})p(\boldsymbol{\tau})p(\boldsymbol{\vartheta})$ is the joint prior distribution. In the high-dimensional setting, computing the posterior distribution is challenging because of the intractable form of the marginal distribution $p(\mathbf{W}; \mathbf{X}, \mathbf{t})$ and the non-conjugate priors on the model parameters. Markov Chain Monte Carlo (MCMC) sampling provides a helpful paradigm for obtaining the required posterior distribution in the Bayesian framework. However, MCMC lacks computational efficiency in large/high-dimensional problems such as VI-MIDAS. Hence, we use the framework of mean-field VI [Jordan et al., 1999, Wainwright et al., 2008, Blei et al., 2017] and approximate the posterior with a variational posterior distribution of the latent variable $\boldsymbol{\ell}$.

We let $q(\boldsymbol{\ell}; \boldsymbol{\nu})$ be the variational posterior distribution with parameter $\boldsymbol{\nu}$. VI minimizes the Kullback-Leibler (KL) divergence,

$$\min_{\boldsymbol{\nu}} \text{ KL}(q(\boldsymbol{\ell}; \boldsymbol{\nu}) \,||\, p(\boldsymbol{\ell}; \mathbf{W}, \mathbf{X}, \mathbf{t}))$$

such that $\text{supp}(q(\boldsymbol{\ell}; \boldsymbol{\nu})) \subseteq \text{supp}(p(\boldsymbol{\ell}; \mathbf{W}, \mathbf{X}, \mathbf{t}))$. On simplification, the optimization problem is equivalent to maximizing the evidence lower bound (ELBO) given by

$$\mathcal{L}(\boldsymbol{\nu}) = \mathbb{E}_{q(\boldsymbol{\ell};\boldsymbol{\nu})}[\log P(\mathbf{W}, \boldsymbol{\ell}; \mathbf{X}, \mathbf{t})] - \mathbb{E}_{q(\boldsymbol{\ell};\boldsymbol{\nu})}[\log q(\boldsymbol{\ell}; \boldsymbol{\nu})], \tag{2}$$

a lower bound on the logarithm of the joint probability of the observations, i.e., $\log P(\mathbf{W}; \mathbf{X}, \mathbf{t})$ [Jordan et al., 1999]. One can further simplify by replacing the joint distribution $P(\mathbf{W}, \boldsymbol{\ell}; \mathbf{X}, \mathbf{t})$ with a product of likelihood and prior distribution $P(\mathbf{W}, \boldsymbol{\ell}; \mathbf{X}, \mathbf{t}) = P(\mathbf{W}; \boldsymbol{\ell}, \mathbf{X}, \mathbf{t})P(\boldsymbol{\ell})$.

Coordinate ascent variational inference provides an efficient framework for approximating the variational posterior in a generative model with conjugate priors satisfying support-matching constraints [Blei et al., 2017]. For a general scenario, such as VI-MIDAS, we transform the support of the latent variable $\boldsymbol{\ell}$ to a real coordinate space using a one-to-one differentiable function

$$\mathbf{T} : \text{supp}(p(\boldsymbol{\ell})) = \mathbb{R}^l \tag{3}$$

and express the transformed variable as $\boldsymbol{\zeta} = \mathbf{T}(\boldsymbol{\ell})$ where $\boldsymbol{\zeta} \in \mathbb{R}^l$. For example, given any latent variable $a \in \mathbb{R}^+$, then for $T = \log$, we have $\zeta = T(a) \in \mathbb{R}$. Similarly, for any $a \in (0, 1)$, we apply the logit transform and write $\zeta = T(a) = \text{logit}(a) = \log(\frac{a}{1-a}) \in \mathbb{R}$. Using a standard transformation, we express the joint distribution of the transformed latent variable $\boldsymbol{\zeta}$ as $P(\mathbf{W}, \mathbf{T}^{-1}(\boldsymbol{\zeta}); \mathbf{X}, \mathbf{t})|\det \mathbf{J}_{\mathbf{T}^{-1}}(\boldsymbol{\zeta})|$. For the unconstrained latent variables $\boldsymbol{\zeta}$, we formulate the mean-field variational posterior distribution as

$$q(\boldsymbol{\zeta}; \boldsymbol{\mu}, \boldsymbol{\sigma}) \sim \mathcal{N}(\boldsymbol{\mu}, \boldsymbol{\Sigma}) = \prod_{i=1}^{l} \mathcal{N}(\zeta_i; \mu_i, \sigma_i),$$

where $\boldsymbol{\mu} = [\mu_1, \ldots, \mu_l] \in \mathbb{R}^l$ is the mean parameter and $\boldsymbol{\sigma} = [\sigma_1, \ldots, \sigma_l] \in \mathbb{R}^{+l}$ is the variance parameter. Now, we reformulate the ELBO (2) in terms $\boldsymbol{\zeta}$ as

$$\begin{aligned}\mathcal{L}(\boldsymbol{\mu}, \boldsymbol{\sigma}) &= \mathbb{E}_{q(\boldsymbol{\zeta};\boldsymbol{\mu},\boldsymbol{\sigma})}[\log P(\mathbf{W}, \mathbf{T}^{-1}(\boldsymbol{\zeta}); \mathbf{X}, \mathbf{t}) + \log|\det \mathbf{J}_{\mathbf{T}^{-1}}(\boldsymbol{\zeta})|] - \mathbb{E}_{q(\boldsymbol{\zeta};\boldsymbol{\mu},\boldsymbol{\sigma})}[\log q(\boldsymbol{\zeta}; \boldsymbol{\kappa})] \\ &= \mathbb{E}_{q(\boldsymbol{\zeta};\boldsymbol{\mu},\boldsymbol{\sigma})}[\log P(\mathbf{W}, \mathbf{T}^{-1}(\boldsymbol{\zeta}); \mathbf{X}, \mathbf{t}) + \log|\det \mathbf{J}_{\mathbf{T}^{-1}}(\boldsymbol{\zeta})|] + \sum_i \log \sigma_i + const. \end{aligned} \tag{4}$$

Here, Gaussian variational distribution is related to the second-order approximation of the posterior around the maximum-a-posteriori (MAP) estimate. In terms of the original parameter $\boldsymbol{\ell}$, variational distribution is non-Gaussian because of the transform $\mathbf{T}$ and its Jacobian. To estimate parameters of the variational posterior after reparameterization, VI-MIDAS solves

$$\widehat{\boldsymbol{\mu}},\ \widehat{\boldsymbol{\sigma}} \equiv \arg\max_{\boldsymbol{\mu},\boldsymbol{\sigma}} \mathcal{L}(\boldsymbol{\mu},\boldsymbol{\sigma}), \tag{5}$$

using the coordinate ascent approach. We solve the optimization problem using stochastic gradient ascent (SGA), which uses automatic differentiation (AD) to compute the gradient and Monte Carlo integration to approximate the expectation [Blei et al., 2017]. AD is applicable when gradient operation is inside the expectation. The estimation approach achieves this by applying an additional elliptical transformation given by $\kappa_i = (\zeta_i - \mu_i)/\exp(v_i)$ where $v_i = \log\sigma_i$. We denote the set of new latent variable as $\boldsymbol{\kappa} = [\kappa_1, \ldots, \kappa_l]$, and reparameterize ELBO as

$$\mathcal{L}(\boldsymbol{\mu},\mathbf{v}) = \mathbb{E}_{q(\boldsymbol{\kappa};\mathbf{0},\mathbf{1})}[\log P(\mathbf{W}, \mathbf{T}^{-1}(\mathbf{S}(\boldsymbol{\kappa})); \mathbf{X}, \mathbf{t}) + \log|\det \mathbf{J}_{\mathbf{T}^{-1}}(\mathbf{S}(\boldsymbol{\kappa}))|] + \sum_i v_i, \tag{6}$$

where $\boldsymbol{\zeta} = \mathbf{S}(\boldsymbol{\kappa}) = \text{diag}[exp(\mathbf{v})]\boldsymbol{\kappa} + \boldsymbol{\mu}$ and $\mathbf{v} = [v_1, \ldots, v_l]$. To execute gradient ascent, we need to compute $\frac{d\mathcal{L}(\boldsymbol{\mu},\mathbf{v})}{d\boldsymbol{\mu}}$ and $\frac{d\mathcal{L}(\boldsymbol{\mu},\mathbf{v})}{d\mathbf{v}}$. Let us represent the derivative of the random variable function inside the expectation as $\frac{da}{d\boldsymbol{\kappa}} = \nabla_{\boldsymbol{\theta}} \log P(\mathbf{W}, \boldsymbol{\theta}; \mathbf{X}, \mathbf{t})\nabla_{\boldsymbol{\kappa}}\mathbf{T}^{-1}(\mathbf{S}(\boldsymbol{\kappa})) + \nabla_{\boldsymbol{\kappa}} \log|\det \mathbf{J}_{\mathbf{T}^{-1}}(\mathbf{S}(\boldsymbol{\kappa}))|$. Then, with reparameterized ELBO, the gradient with respect to $\boldsymbol{\mu}$ and $\mathbf{v}$ is given by

$$\frac{d\mathcal{L}(\boldsymbol{\mu},\mathbf{v})}{d\boldsymbol{\mu}} = \mathbb{E}_{q(\boldsymbol{\kappa};\mathbf{0},\mathbf{1})}\left[\frac{da}{d\boldsymbol{\kappa}}\right] \quad \text{and} \quad \frac{d\mathcal{L}(\boldsymbol{\mu},\mathbf{v})}{d\mathbf{v}} = \mathbb{E}_{q(\boldsymbol{\kappa};\mathbf{0},\mathbf{1})}\left[\frac{da}{d\boldsymbol{\kappa}} \odot exp(\mathbf{v}) \odot \boldsymbol{\kappa}\right] + \mathbf{1}. \tag{7}$$

We compute the gradient inside the expectation using automatic differentiation and then approximate the expectation using MC integration by drawing $m$ (typically $m = 1$) from the standard normal distribution. This step results in a noisy and unbiased estimate of the gradient of the ELBO. Using the gradient estimate, we update the variational parameters $\{\boldsymbol{\mu}, \mathbf{v}\}$ via stochastic optimization given by

$$\boldsymbol{\mu}^{(i+1)} \leftarrow \boldsymbol{\mu}^{(i)} + \boldsymbol{\xi}_{\boldsymbol{\mu}}^{(i)} \odot \frac{d\mathcal{L}(\boldsymbol{\mu},\mathbf{v})}{d\boldsymbol{\mu}}, \qquad \text{and} \qquad \mathbf{v}^{(i+1)} \leftarrow \mathbf{v}^{(i)} + \boldsymbol{\xi}_{\mathbf{v}}^{(i)} \odot \frac{d\mathcal{L}(\boldsymbol{\mu},\mathbf{v})}{d\mathbf{v}}$$

where $\boldsymbol{\xi}_{\boldsymbol{\mu}}^{(i)}$ and $\boldsymbol{\xi}_{\mathbf{v}}^{(i)}$ are step-size. The computation procedure is guranteed to converge to a local maximum of the ELBO under the condition of a sufficiently decaying step-size sequence. In particular, we use an adaptive step-size sequence [Duchi et al., 2011] to meet this condition given by

$$\boldsymbol{\xi}_{\boldsymbol{\mu}}^{(i)} = \vartheta \times i^{-1/2+\varepsilon} \times \left(\varsigma + \sqrt{\boldsymbol{\varrho}_{\boldsymbol{\mu}}^{(i)}}\right)^{-1} \quad \text{and} \quad \boldsymbol{\xi}_{\mathbf{v}}^{(i)} = \vartheta \times i^{-1/2+\varepsilon} \times \left(\varsigma + \sqrt{\boldsymbol{\varrho}_{\mathbf{v}}^{(i)}}\right)^{-1}, \tag{8}$$

where $\vartheta$ is the learning rate, $\boldsymbol{\varrho}_{\boldsymbol{\mu}}^{(i)} = \alpha\frac{d\mathcal{L}(\boldsymbol{\mu},\mathbf{v})}{d\boldsymbol{\mu}} + (1-\alpha)\boldsymbol{\varrho}_{\boldsymbol{\mu}}^{(i-1)}$ and $\boldsymbol{\varrho}_{\mathbf{v}}^{(i)} = \alpha\frac{d\mathcal{L}(\boldsymbol{\mu},\mathbf{v})}{d\mathbf{v}} + (1-\alpha)\boldsymbol{\varrho}_{\mathbf{v}}^{(i-1)}$ is the curvature. In practice, we search for the optimal learning rate such that $\vartheta \in \{0.01, 0.1, 1, 10, 100\}$. Other parameters, such as $\varepsilon = 10^{-16}$, $\varsigma = 1$ are chosen to prevent zero division, and $\alpha = 0.1$ are chosen in order to give more weight to the past curvature. This suggested approach to parameter estimation comes under the framework of Automatic Differentiation Variational Inference (ADVI) [Kucukelbir et al., 2017]. The approach is implemented in the probabilistic programming language Stan [Carpenter et al., 2017]. We have summarized the parameter estimation procedure in Algorithm 1.

**Algorithm 1** Automatic Differentiation Variational Inference for VI-MIDAS

**Given**: Count abundance data $\mathbf{W}$, covariates $\mathbf{X}$, province $\mathbf{P}$, ocean depth $\mathbf{D}$, model $P(\mathbf{W}, \mathbf{T}^{-1}(\mathbf{S}(\boldsymbol{\kappa})); \mathbf{X}, \mathbf{t})$ where $\mathbf{S}(\boldsymbol{\kappa}) = \text{diag}[exp(\mathbf{v})]\boldsymbol{\kappa} + \boldsymbol{\mu}$.

Initialize variational posterior parameter $\boldsymbol{\mu}^{(1)} = \mathbf{0}, \mathbf{v}^{(1)} = \mathbf{0}$, $i = 1$.

Search for learning parameter rate $\vartheta$ over a set of finite values.

**repeat**

  Draw $m = 1$ sample from $\mathcal{N}(\mathbf{0}, \mathbf{I})$.

  Estimate noisy gradient $\frac{d\mathcal{L}(\boldsymbol{\mu},\mathbf{v})}{d\boldsymbol{\mu}}$ and $\frac{d\mathcal{L}(\boldsymbol{\mu},\mathbf{v})}{d\mathbf{v}}$ using MC integration; see equation (7).

  Calculate step-size $\boldsymbol{\xi}_{\mu}^{(i)}$ and $\boldsymbol{\xi}_{\mathbf{v}}^{(i)}$ using ADAM; see equation (8).

  Update $\boldsymbol{\mu}^{(i+1)} \leftarrow \boldsymbol{\mu}^{(i)} + \boldsymbol{\xi}_{\mu}^{(i)} \odot \frac{d\mathcal{L}(\boldsymbol{\mu},\mathbf{v})}{d\boldsymbol{\mu}}$ and $\mathbf{v}^{(i+1)} \leftarrow \mathbf{v}^{(i)} + \boldsymbol{\xi}_{\mathbf{v}}^{(i)} \odot \frac{d\mathcal{L}(\boldsymbol{\mu},\mathbf{v})}{d\mathbf{v}}$.

  $i \leftarrow i + 1$

**until** $|\mathcal{L}(\boldsymbol{\mu}^{(i+1)}, \mathbf{v}^{(i+1)}) - \mathcal{L}(\boldsymbol{\mu}^{(i)}, \mathbf{v}^{(i)})| < \epsilon$ where $\epsilon = 0.01$

**return** $\boldsymbol{\mu}^* = \boldsymbol{\mu}^{(i)}, \mathbf{v}^* = \mathbf{v}^{(i)}$.

#### 3.3 Hyperparameter tuning

We estimate the model parameters using the ADVI approach implemented in the probabilistic programming language Stan. Please see Appendix **??** for the Stan code implementing the generative model for the microbial abundance data. VI-MIDAS requires specifying the dimension $k$ of the latent variable and the hyper-parameters $\{\lambda, \upsilon\}$ of the sparsity-inducing Laplace priors and the inverse-Cauchy prior of the generative model. In the hyperparameter tuning step, the procedure selects 50 random settings from a range of possible values of the parameters given by $k \in \{10, 16, 30, 50, 80, 100, 150, 200, 500\}$, $\lambda \in (0.01, 3000)$ and $\upsilon \in (0.03125, 0.5)$. To evaluate these settings, we split the data into five folds with 90% training and 10% testing. We use Algorithm 1 to estimate parameters on the former and then evaluate the estimate on the latter using out-of-sample log pointwise predictive density (LLPD) [Gelman et al., 2013, Blei et al., 2017]. Figure S6 reports the test sample LLPD to evaluate the model fit for the selected settings. We select all the settings in the range of one standard deviation of the most significant value of the LLPD; see Table S2 in the supplementary materials.

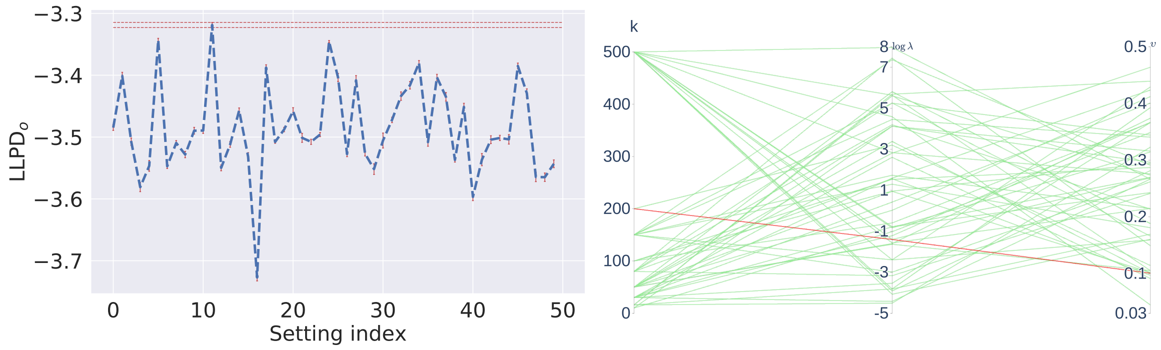


(a) (b)

Figure S6: Hyperparameter tuning: a) out-of-sample log pointwise predictive density (LLPD) comparing fifty different settings randomly selected from a range possible values: taxa-specific feature vector length $k \in \{10, 16, 30, 50, 80, 100, 150, 200, 500\}$, hyperparameter for Laplace priors $\lambda \in (0.01, 3000)$ and the hyperparameter for inverse-Cauchy priors $\upsilon \in (0.03125, 0.5)$; b) parallel coordinate plot highlighting the hyperparameter settings of high (H : dark red) and low (L: light green) mean values of LLPD. The analysis has selected $k = 200$, $\lambda = 0.246$ and $\upsilon = 0.10063$ as the values of the latent variable dimension and the hyperparameters with the highest LLPD $= -3.32$.

##### 3.4 Ablation Study

The analysis quantifies the relative importance of each of the components, i.e., interaction(-I), biogeochemical environment(-E), province(-P), ocean depth(-D), and seasonality(-S) in the VI-MIDAS model. For each component in the VI-MIDAS model, we consider a component-excluded model. For instance, the model that excludes the interaction component is denoted VI-MIDAS(-I). Similarly, VI-MIDAS(-E), VI-MIDAS(-P), VI-MIDAS(-D) and VI-MIDAS(-S) denote the models excluding the geochemical environment(E), province(P), ocean depth(D) and seasonality(S) components, respectively. For the selected setting from the hyperparameter tuning step, we evaluate each component-excluded model in terms of the out-of-sample LLPD. From the comparison results in Table S4, we observe that the seasonality (S) is least important and the ocean depth(D) component is most important.

Table S4: Out-of-sample log-likelihood posterior predictive density (LLPD) of the full model and after ablation of the environmental(E), province(P), ocean depth(D), seasonality(S), and latent interaction (I) component.

| **Model** | VI-MIDAS | VI-MIDAS(-E) | VI-MIDAS(-P) | VI-MIDAS(-D) | VI-MIDAS(-S) | VI-MIDAS(-I) |
|---|---|---|---|---|---|---|
| **LLPD** | -3.3**322** | -3.3**554** | -3.3**398** | -3.3**882** | -3.3**335** | -3.3**377** |

##### 3.5 Model sensitivity analysis

The objective function maximizing the ELBO is non-convex; hence, the estimates of the parameters are sensitive to their initial values. For each of the settings selected in Table S2, we estimate the model parameters on complete data for fifty random initializations. The parameters estimated from the random initializations are evaluated based on the fitted LLPD [Gelman et al., 2013]. Out of fifty different parameter estimates for the selected setting, we select the one with the most significant value of fitted LLPD. Based on the variational posterior parameter estimate, we generate 100 posterior samples of the latent variables $\boldsymbol{\ell}$.

##### 3.6 Model fit diagnostic

After performing the model sensitivity analysis, we obtain the estimate of the VI-MIDAS model parameters $\boldsymbol{\ell}$. The model fit is numerically evaluated using the posterior predictive check [Rubin, 1984, Gelman et al., 2013] on the full data. The procedure requires generating $m = 100$ posterior samples, denoted by the random variable $\mathbf{W}^{rep} = [w_{ij}^{rep}] \in \mathbb{R}_+^{n \times q}$, and then computing the p-value of the model fit as

$$\text{p-value} = p(t(\mathbf{W}^{rep}) < t(\mathbf{W})),$$

where $t$ is the test statistic. In practice, we use the test statistics $t(\mathbf{W}^{rep}) = \mathbf{E}(\log p(\mathbf{W}^{rep}|\boldsymbol{\ell}))$ and $t(\mathbf{W}) = \mathbf{E}(\log p(\mathbf{W}|\boldsymbol{\ell}))$. For the selected setting, we have p-value $= 0.53$, and thus we fail to reject the hypothesis that the posterior samples are different from the fitted $\mathbf{W}$. A simpler test statistic is $t(\mathbf{w}) = \mathbf{w}$. In this case too, we have p-value $= 0.59$. We visually examine the model fit by comparing the abundance data with its predicted value and the error; see Figure S7(a-c). Figure S7 (d) reports the convergence of the ELBO using ADVI. Finally, we compare the observed and estimated abundance profiles of the species using the Q-Q plot (for distribution) and the scatter plot; see Figure S7(e-f).

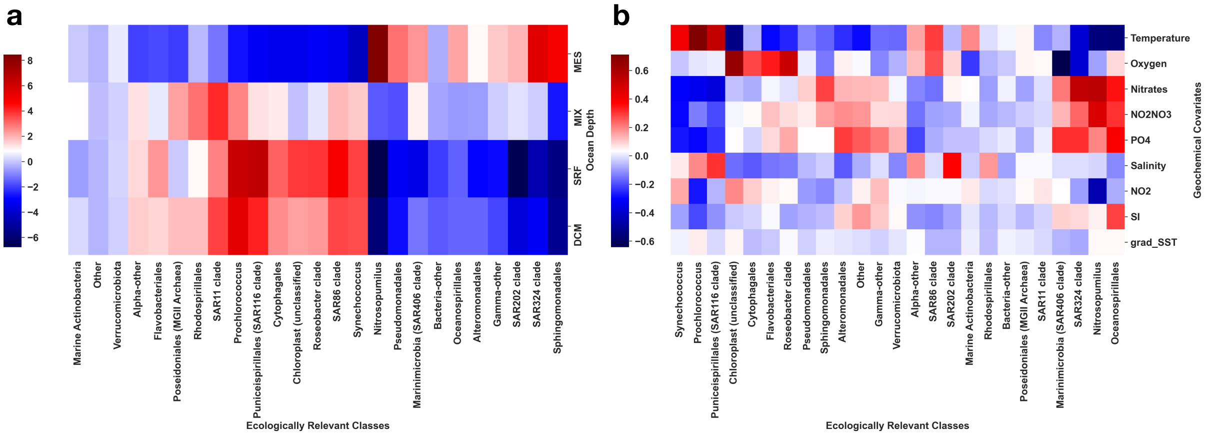


Figure S7: Model fit diagnostic: a) Heatmap showing the abundance profile $\log(\mathbf{W}+1)$ of 1378 species at $n = 139$ distinct geographical locations across the globe; b) Expected value of the abundance using the hyperparameter corresponding to best model fit; c) Error plot representing the absolute value of the difference between the observed and expected species abundance, denoted by $|\log \mathrm{E}(W) - \log \mathbf{W}|$; d) Convergence of the ELBO with conformable rank $k = 16$ and hyperparameter $\lambda = 398.199$ and $\upsilon = 0.04904$; e) Q-Q plot comparing the observed and estimated abundance profile of the species; f) Scatter plot comparing entries of the CLR transform of the observed and estimated microbial species abundance, given by $\log \mathbf{W}/\mathbf{T}$ and $\log \mathrm{E}(\mathbf{W})/\mathbf{T}$, respectively; g) Histogram comparing the distribution of the observed and expected species abundance in terms of $\log \mathbf{W}$ and $\log \mathrm{E}(\mathbf{W})$.

### 4 Network analysis of mutualistic and competitive interaction

In the model inference, we have considered the top five positive and top five negative entries in the rows of $\mathbf{I}$ to learn about each taxa' most significant mutualistic and competitive interactions. Based on the significant entries in $\mathbf{I}$, we separately define the adjacency matrix of the mutualistic (positive entries) and competitive (negative entries) interactions and then visualize them through the network plots; see Figure S8 (a-b).

We have also identified the most important OTUs/taxa in the interaction networks (based on adjacency matrix; see Figure S8 (a-b)) as a hub (denoted with a "*" shape) with a degree greater than one hundred. The associations of each of the hub nodes with other taxa (in term ERC types) are summarized in the mutualistic and competitive interaction heatmap in the Figure S8 (c). Each of the entries in the heatmap reports the fraction of an ERC type (x-axis) associated with a hub node (y-axis). We have further summarized the most important associations (greater than 0.5 in Figure S8) of hub nodes in the mutualistic and competitive interactions network in Table S5.

The set of OTUs (out of a total of 1378) that exhibit the most significant mutualistic interaction includes {OTU2, OTU3, OTU4, OTU11, OTU13, OTU18, OTU25, OTU21, OTU32, OTU36, OTU48, OTU90}; see Table S5 for the details of taxonomic rank. The ECR types of their hub nodes mainly comprise SAR11 clade, Synechococcus, Prochlorococcus, Rhodospirillales, and Planctomycetes. Based on the ERC type of the nodes connected to OTU2, OTU4, OTU25 and OTU90 (see Table S5), we deduce the existence of subtypes of Rhodospirillales that may differ in ecology and metabolic functions. Using organic substrates as the carbon source, Rhodospirillales can grow in a variety of conditions such as a) anaerobically in the light, b) aerobically in the dark, and c) fermentatively (anaerobically) in the dark [Alber, 2009]. Similarly, based on the nodes connected to OTU11, OTU13, OTU18, OTU22 and OTU32, one can learn about the metabolic versatility of the most abundant ERC type SAR11 clade; and its ability to exhibit mutualistic relationships with other species.

Now, the set of OTUs that exhibit the most significant competitive interactions includes {OTU19, OTU37, OTU354, OTU1139, OTU1563, OTU1824, OTU2578, OTU5312, OTU4327, OTU4683, OTU4704}; see Table S5 for the details of taxonomic rank. In competitive interactions, one species may discourage the abundance of the other either directly (gazing) or indirectly (metabolic pathway). Most of the OTU that act as hub nodes have very low mean abundance except {OTU19, OTU37, OTU354}. Based on the competitive interactions of OTU19, the analysis suggests that SAR86 clade discourages the abundance of taxa with the ERC indicators Bacteria-other, Oceanospirillales, SAR202 clade, SAR324 clade, Pseudomonadales, Nitrosopumilus, and Sphingomonadales. Similarly, OTU19, a Gamma-other bacteria, reduces the abundance of Poseidoniales (MGII Archaea) and Prochlorococcus, and OTU354, a Marine Actinobacteria, reduces the abundance of Pseudomonadales and Sphingomonadales.

Table S5: Description of the hub nodes in mutualistic and competitive interactions network (see Figure S8 (a-b) of the SM) in terms of their ID, ERC indicator, and mean abundance and their most significant associations (greater than 0.5 in Figure S8(c) of the SM) with other ERC types.

| Significant Interaction Type | HUB Marker | OTU ID | HUB ERC | Abundance (Mean) | Connection ERC |
|---|---|---|---|---|---|
| | 4 | OTU2 | Rhodospirillales | 227.3 | Rhodospirillales, Alpha-other, SAR86 clade, SAR11 clade, Synechococcus, Puniceispirillales (SAR116 clade), Prochlorococcus |
| | 5 | OTU3 | Prochlorococcus | 193.7 | Poseidoniales (MGII Archaea), SAR86 clade, Synechococcus, Prochlorococcus, Puniceispirillales (SAR116 clade) |
| | 12 | OTU4 | Rhodospirillales | 177.7 | Gamma-other, SAR202 clade, Marinimicrobia (SAR406 clade), SAR324 clade, Nitrosopumilus |
| | 2 | OTU11 | SAR11 clade | 99.1 | SAR11 clade |
| | 11 | OTU13 | SAR11 clade | 97.7 | SAR202 clade |
| | 1 | OTU18 | SAR11 clade | 90.0 | Rhodospirillales, SAR86 clade, Synechococcus, SAR11 clade, Prochlorococcus, Puniceispirillales (SAR116 clade) |
| **Mutualistic** | 0 | OTU25 | Rhodospirillales | 86.5 | Poseidoniales (MGII Archaea), SAR202 clade, Synechococcus, Prochlorococcus |
| | 14 | OTU21 | SAR11 clade | 85.2 | Nitrosopumilus, SAR324 clade |
| | 3 | OTU32 | SAR11 clade | 73.1 | Poseidoniales (MGII Archaea) |
| | 10 | OTU36 | SAR86 clade | 73.0 | Puniceispirillales (SAR116 clade) |
| | 13 | OTU48 | Planctomycetota | 72.0 | SAR202 clade, SAR324 clade |
| | 6 | OTU90 | Rhodospirillales | 51.2 | Alteromonadales, Gamma-other, Pseudomonadales, SAR202 clade |
| | 16 | OTU19 | SAR86 clade | 80.7 | Bacteria-other, Oceanospirillales, SAR202 clade, SAR324 clade, Pseudomonadales, Nitrosopumilus, Sphingomonadales |
| | 2 | OTU37 | Gamma-other | 71.3 | Poseidoniales (MGII Archaea), Prochlorococcus |
| | 9 | OTU354 | Marine Actinobacteria | 22.1 | Pseudomonadales, Sphingomonadales |
| | 15 | OTU1139 | Flavobacteriales | 9.0 | Alteromonadales, Oceanospirillales, SAR202 clade, SAR324 clade, Nitrosopumilus, Pseudomonadales, Sphingomonadales |
| | 1 | OTU1563 | Flavobacteriales | 8.2 | Poseidoniales (MGII Archaea), SAR11 clade, SAR86 clade, Synechococcus, Prochlorococcus |
| **Competitive** | 0 | OTU1824 | Oceanospirillales | 6.9 | Alpha-other, Marine Actinobacteria, Rhodospirillales, Chloroplast (unclassified), Cytophagales, Puniceispirillales (SAR116 clade), SAR11 clade, SAR86 clade, Poseidoniales (MGII Archaea), Synechococcus, Prochlorococcus |
| | 13 | OTU2578 | Marine Actinobacteria | 6.0 | SAR202 clade, SAR324 clade, Nitrosopumilus, Pseudomonadales, Sphingomonadales |
| | 3 | OTU5312 | Chloroplast (unclassified) | 4.6 | Prochlorococcus |
| | 11 | OTU4327 | Cytophagales | 4.6 | Bacteria-other, Roseobacter clade |
| | 4 | OTU4683 | Alteromonadales | 4.5 | Prochlorococcus, Chloroplast (unclassified), Puniceispirillales (SAR116 clade), Synechococcus |
| | 5 | OTU4704 | Alteromonadales | 4.5 | SAR202 clade, SAR324 clade, Nitrosopumilus, Sphingomonadales, Pseudomonadales |

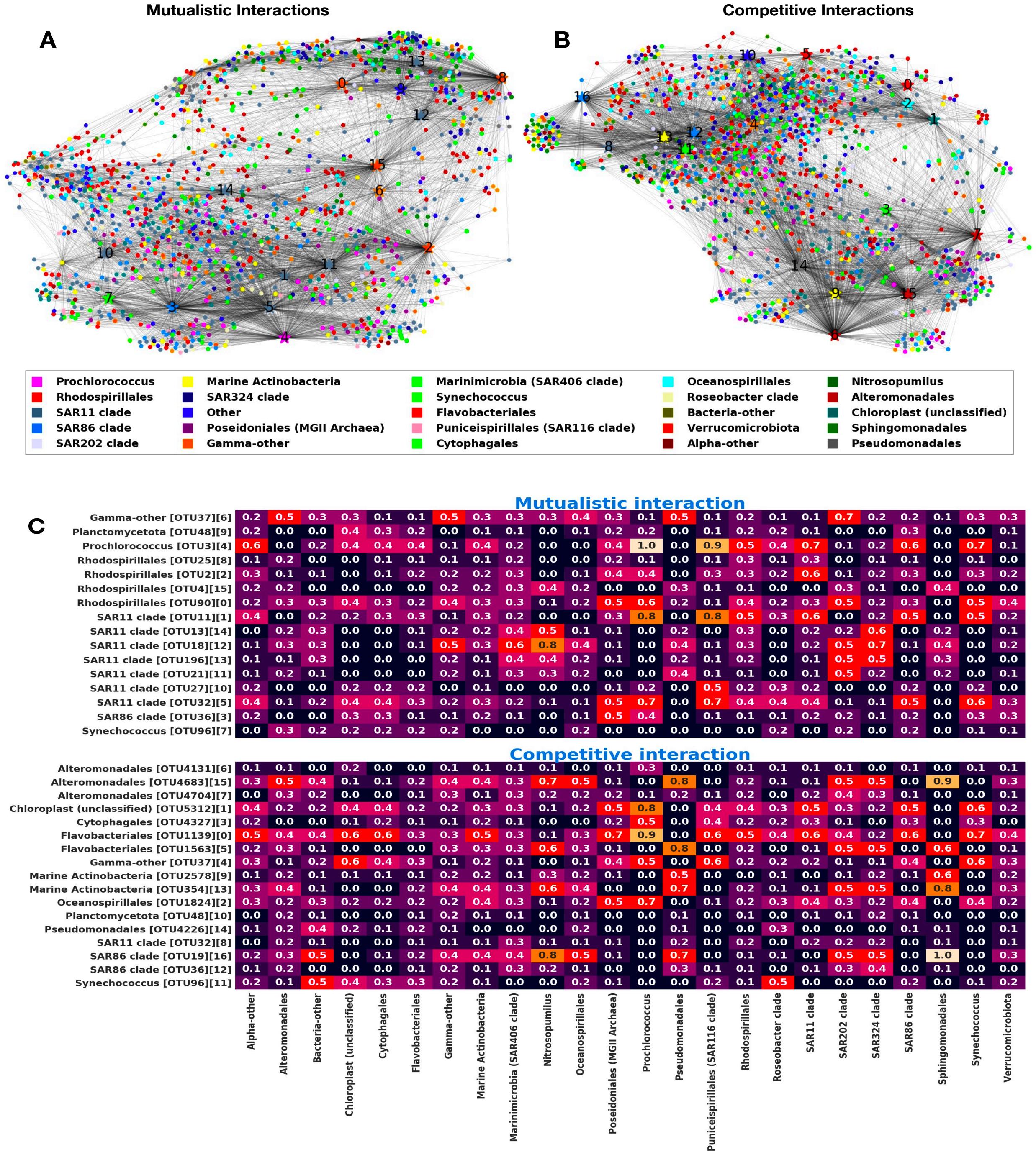


Figure S8: Interactions among species: Based on the significant entries in **I**, we separately define the adjacency matrix of the mutualistic (positive entries) and competitive (negative entries) interactions and then visualize them through the network plots. a) Network plot based on the top 5 positive interactions of each of the OTUs with "*" denoting the ten hub nodes identified with degree greater than 100. b) Network plot based on the top 5 negative interactions of each of the OTUs with "*" representing the 11 hub nodes identified with degree greater than 100. c) Distribution of the association of the hub nodes (y-axis) with other species. Entries in the heatmap show association of a hub node with the fraction of each of the ERC types (x-axis).

Roberto de la Cruz and Jan-Ulrich Kreft. Geometric mean extension for data sets with zeros. *arXiv preprint arXiv:1806.06403*, 2018.

John Duchi, Elad Hazan, and Yoram Singer. Adaptive subgradient methods for online learning and stochastic optimization. *Journal of machine learning research*, 12(Jul):2121–2159, 2011.

AB Gelman, JB Carlin, HS Stern, DB Dunson, A Vehtari, and D Rubin. Bayesian data analysis third edition. boca raton. *FL: CRC Press.[Google Scholar]*, 2013.

Michael I Jordan, Zoubin Ghahramani, Tommi S Jaakkola, and Lawrence K Saul. An introduction to variational methods for graphical models. *Machine learning*, 37(2):183–233, 1999.

Alp Kucukelbir, Dustin Tran, Rajesh Ranganath, Andrew Gelman, and David M Blei. Automatic differentiation variational inference. *The Journal of Machine Learning Research*, 18(1):430–474, 2017.

Aditya Mishra et al. Robust regression with compositional covariates. *arXiv preprint arXiv:1909.04990*, 2019.

Donald B Rubin. Bayesianly justifiable and relevant frequency calculations for the applied statistician. *The Annals of Statistics*, pages 1151–1172, 1984.

Pixu Shi, Anru Zhang, Hongzhe Li, and Others. Regression analysis for microbiome compositional data. *Ann. Appl. Stat.*, 10(2):1019–1040, 2016.

Martin J Wainwright, Michael I Jordan, et al. Graphical models, exponential families, and variational inference. *Foundations and Trends® in Machine Learning*, 1(1–2):1–305, 2008.

Xinyan Zhang, Himel Mallick, Zaixiang Tang, Lei Zhang, Xiangqin Cui, Andrew K Benson, and Nengjun Yi. Negative binomial mixed models for analyzing microbiome count data. *BMC bioinformatics*, 18(1):4, 2017.
